# RomX, a novel prokaryotic regulator, links the response receiver domain of RomR with GTP-bound MglA for establishing *Myxococcus xanthus* polarity

**DOI:** 10.1101/2024.02.20.581209

**Authors:** Sukanya Chakraborty, Pananghat Gayathri

## Abstract

Cell polarity specification and reversals are distinctive features of motility of the soil bacterium *Myxococcus xanthus*. The bacterial small Ras-like GTPase, MglA, serves as a key player orchestrating these polarity oscillations. RomR, a response regulator, along with its partner RomX, has been identified to function as a guanine nucleotide exchange factor (GEF) for MglA, crucial for its polar recruitment. In this study, we determine the crystal structure of RomX, a protein of a hitherto unknown fold. RomX consists of a three-helix bundle, identified to be the same fold as the stalk domain of atlastin, a member of the dynamin family of GTPases. From our structure-based sequence analysis for proteins of a similar fold, we observe the co-occurrence of the RomX fold with response receiver domains in several bacterial response regulators. Based on mutational analysis and affinity measurements, we conclude that the helix-1 of RomX mediates the interaction with MglA-GTP, while helix-3 of RomX interacts with the RomR N-terminal receiver (REC) domain. We demonstrate that the binding between MglA and RomX is exclusively in the presence of GTP. The absence of additional stimulation of RomX GEF activity in the presence of RomR-REC supports the mutually exclusive interface on RomX for RomR and MglA interaction. Collectively, our findings validate the positioning of RomX between MglA and RomR-REC, providing insights into the concerted action of the RomR/RomX complex at both poles in driving MglA localization within polarized cells.

## Introduction

Rod-shaped *Myxococcus* cells possess a specified front-rear polarity that governs their directional movement^1–3^. The core of this oscillatory system consists of proteins exhibiting distinct subcellular localization patterns that determine the front-rear cell polarity^4,5^. Interestingly, *Myxococcus xanthus* undergoes frequent reversals in cell polarity, a phenomenon important in shaping its unique motility pattern which is also crucial for sustaining its life cycle^6–8^. This polarity reversal, associated with the redistribution of polarly localized proteins^9–11^, prompts the cell to travel in the opposite direction. The frequency of cell polarity reversals is finely tuned by external chemosensory cues, which orchestrate the dynamic behaviour of *Myxococcus xanthus* with respect to nutrient availability^12,13^.

MglA is a prokaryotic small Ras-like GTPase, which acts as the major regulator driving *Myxococcus xanthus* polarity reversals^14–16^. Similar to other well-characterized eukaryotic small Ras-like GTPases, it switches between the active GTP-bound and inactive GDP-bound states^17–19^. This is associated with conformational change at regions near the nucleotide-binding pocket, also referred to as the Switch regions, that affects its localization and interaction with other proteins driving motility and reversals^17,20,21^. The active GTP bound state localizes to the leading cell pole and relocates to the opposite pole when the reversal switch is triggered^17,18^. When the bound GTP is hydrolyzed, the protein in the GDP-bound state detaches from the pole and diffuses in the cytoplasm. In the active GTP bound state, it interacts with SgmX, an effector, and also plays a role in the correct polar localization of PilB and PilT ATPases, all of which are critical to S-motility^22–24^. It also interacts with AglZ in the GTP state, localizing to the focal adhesion complex-like gliding motors, and manifesting cell movement^15,25–28^.

Much like other small Ras-like GTPases, MglA has a very slow intrinsic GTP hydrolysis rate. This results in the requirement of a GTPase activating protein (GAP) which can assist GTP hydrolysis, thereby causing the GTPase to Switch from its “ON” GTP state to the “OFF” GDP state. Further, MglA has a high affinity to GDP over GTP^21^. This necessitates the requirement of a GEF that assists MglA in losing its affinity towards GDP, which consequently helps GTP association. Hence GEFs are essential to reactivate MglA from the “OFF” GDP bound state back to its “ON” GTP bound state. Accordingly, *Myxococcus xanthus* MglA is regulated by various interactors that sense or modulate the bound nucleotide state, together driving the polarity reversal of this bacterium.

MglB acts as the cognate GAP for MglA and accelerates the intrinsic slow GTP hydrolysis activity of MglA which in turn is further accelerated by a co-GAP, RomY^17–21,29^. MglB consists of a Roadblock domain, which has been shown to exert the GAP activity indirectly by orienting MglA active site residues in a conformation favouring GTP hydrolysis^20,21,30^. In addition, MglB possesses a C-terminal extension to the Roadblock domain (MglB-Ct) which was shown to trigger the conformational switch of MglA from GDP to GTP bound states^21,31^. Through an allosteric mechanism, MglB-Ct causes the strongly bound GDP to dissociate and consequently favours GTP binding, thereby acting as a candidate GEF for MglA^21,31^. Simultaneous to the report of the GEF activity of MglB, a new regulator in the polarity reversal module, namely RomX was identified to act as a GEF for MglA together with its interacting partner RomR^32^. RomR is a multi-domain protein with an N-terminal response receiver domain (REC; 1-116 residues) and a C-terminal output domain (117-420)^33^. The output domain consists of a long-disordered region followed by a conserved C-terminal tail (Ct region; residues 369-420). Deletion of the RomR-REC domain impairs cellular reversals^33^. This suggests that the REC domain is essential to trigger RomR relocalization, which is critical to driving reversals^34^. Both RomR and RomX were found to be essential for driving A-motility in *M. xanthus*^32,33^. The C-terminal output domain was critical for the correct polar localization of RomR^33,35^.

RomX co-occurs with its partner RomR across genomes^32^. In *M. xanthus* it was reported that the RomX polar localization is regulated by RomR. RomR interacts with RomX which further interacts with MglA to form a hetero-trimeric complex and helps recruit GTP-bound MglA (MglA-GTP) to the leading pole^32^. RomX was shown to act as the active guanine nucleotide exchange factor (GEF) for MglA^32^. Together with RomR, it was observed to accelerate the GDP to GTP exchange on MglA compared to RomX alone^32^. The main role of the GEF activity of RomR/RomX is hypothesized to be for recruiting MglA-GTP at or near the leading pole of the cell^32^. In addition to its function at the leading pole, RomR/RomX also co-localizes with MglA-GTP in the Agl/Glt focal adhesion complexes driving gliding motility^32^. However, in a polarized cell, RomR/RomX localizes in a bipolar asymmetric cluster, with a bigger cluster at the lagging pole, where MglA has to be excluded^32,33,36^. This leads us to a conundrum of how RomR/RomX could function to maintain the polar localization of MglA as observed in *Myxococcus* motility.

With the goal of obtaining the mechanism of action of RomR/RomX, we report the crystal structure of RomX as a novel alpha-helical protein. Significantly, we observe RomX fold to be associated with the GTPase domain of the dynamin family of GTPases and also often to co-occur with response receiver domains of several bacterial response regulators. A domain-wise dissection of RomR reveals that the receiver domain interacts with RomX. Our detailed biochemical characterization of the interacting interface of MglA and RomX highlights the residues important for complex formation exclusively in the GTP-bound conformation. We demonstrate that RomX is sandwiched between MglA and RomR, and forms the preliminary anchor to selectively recruit MglA-GTP to the leading pole and the focal adhesion complexes. Our observations shed light on how an active mechanism of exclusion of MglA from the lagging pole involves potential active cycling of GEF and GAP activities, regulated through the multi-domain organization of RomR.

## Results

### RomX is monomeric and comprises a three-helix bundle

The hexahistidine-tagged RomX construct was highly soluble upon heterologous expression in *E.coli* and was purified by affinity chromatography. (Supplementary Fig. S1A). However, RomX, despite being an 11.4 kDa protein, eluted aberrantly (not according to the expected monomeric elution volume) at 13.3 ml in analytical size exclusion chromatography (SEC) (Supplementary Fig. S1B). The secondary structure prediction of RomX reveals it to be a predominantly alpha-helical protein (Supplementary Fig. S1C). To distinguish the oligomeric state of RomX, we performed SEC coupled with multi-angle light scattering (MALS) to confirm its oligomeric state. In SEC-MALS, the observed molar mass was 13.6 kDa (Fig. 1A), consistent with the theoretical molecular weight of 11.4 kDa for the monomer. Hence, we concluded that RomX was monomeric in solution. CD spectroscopy provided an alpha helical signature which confirmed that RomX was predominantly composed of alpha helices (Supplementary Fig. S1D).

**Figure 1.**
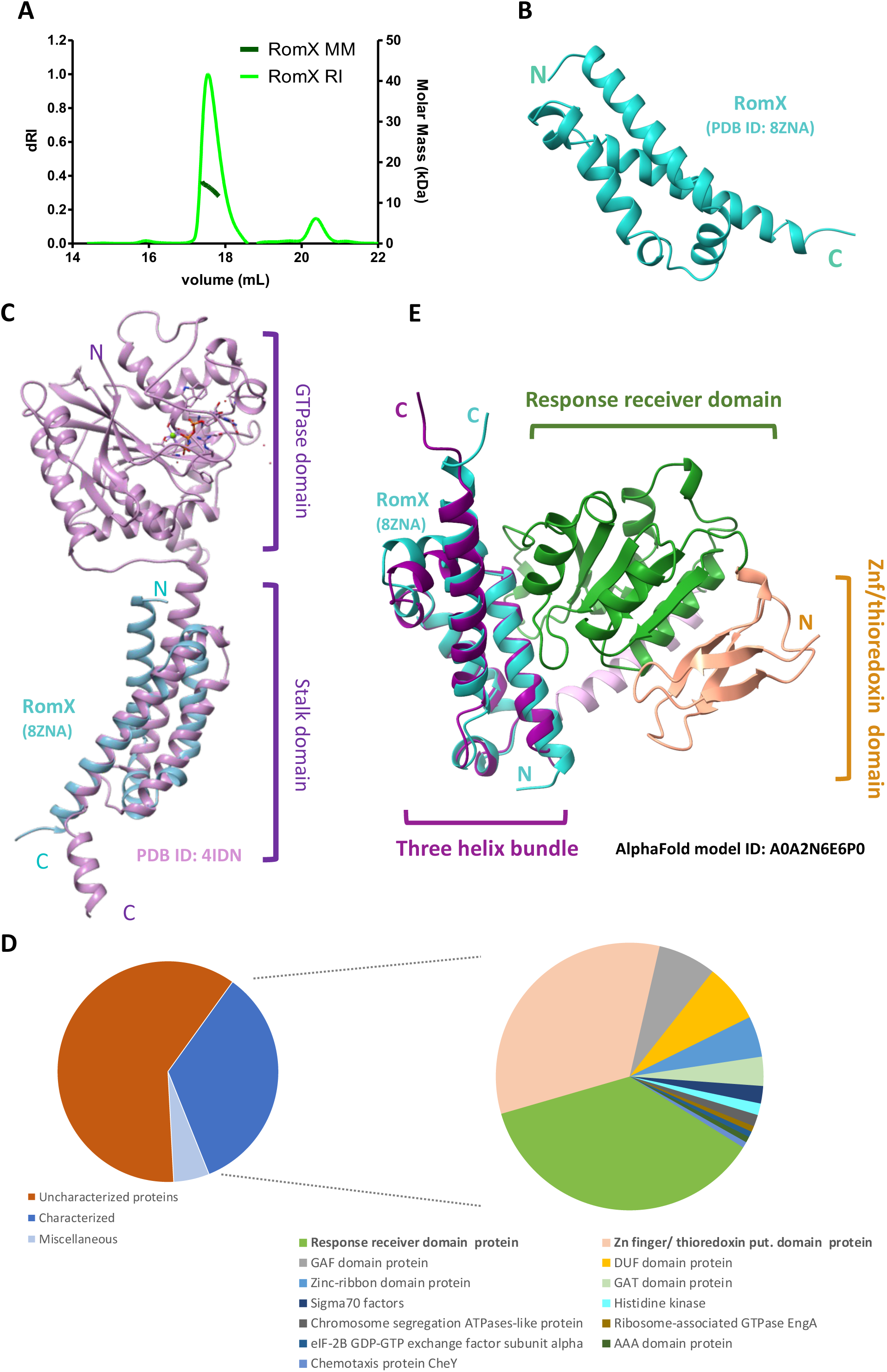
RomX is monomeric, alpha-helical, and consists of a fold associated with GTPases and response regulators. A: SEC-MALS run for RomX using Superdex 200 column. The protein elutes at 17.7 ml and the observed molar mass is 13.6 kDa (±1.964%). The molar mass is represented in dark green and the dRI is represented in light green. B: Crystal structure of C-His RomX solved with anomalous diffraction. There was one molecule in the asymmetric unit. The N and C termini of the protein are labelled. C: Superposition of RomX structure (PDB ID: 8ZNA) on the stalk domain of atlastin (PDB ID: 4IDN), a member of the dynamin family GTPase. D: Pie charts representing the hits from the AlphaFold database using the Foldseek tool. The left plot shows the representation of characterized versus uncharacterized proteins. The proteins of miscellaneous unrelated folds are shown as a subset of characterized proteins. The remainder of the relevant characterized proteins are represented on the right and the respective folds are described in the legend below. E: Superposition of RomX structure (PDB ID: 8ZNA) with a response regulatory domain-containing protein from *Delsulfuromonas sp* (AlphaFold ID: A0A2N6E6P0). The receiver domain and the Zn finger/thioredoxin domains are shown in green and coral respectively. The pink helix in the background is an additional helix between the receiver domain and the RomX-like helical domain. The experimental structure of RomX (PDB ID: 8ZNA, in cyan) is superposed on the helical domain (in dark purple)

Since RomX was of unknown fold with no known homologs from sequence-based searches when this study was initiated, we employed anomalous diffraction from selenomethionine-labeled protein to solve the crystal structure of RomX (Supplementary Fig. S1E). Single-wavelength anomalous scattering data was used to solve the structure to an overall resolution of 1.9 Å for the best crystal (Protein Data Bank [PDB] ID: 8ZNA; Table 1). Upon determining the structure and subsequent refinement, RomX was indeed found to be an alpha-helical protein (Fig. 1B), in which the helices constituted a three-helix bundle. There was one molecule in the asymmetric unit, and crystal-packing analysis further confirmed that it was monomeric.

**Table 1.** Data Collection and refinement statistics of RomX. (Values in parentheses denote the last resolution shell)

|  |  |
| --- | --- |
| Collected at | Diamond Light Source, I03 |
| Wavelength | 0.97625 |
| **Data Collection Statistics** |  |
| Space group | P2<sub>1</sub>2<sub>1</sub>2<sub>1</sub> |
| a,b,c (Å) | 33.32, 56.08, 68.62 |
| $\alpha$, $\beta$, $\gamma$ (°) | 90, 90, 90 |
| Resolution (Å) | 29.95 – 1.89 |
| No. of unique reflections | 266856 (8682) |
| R<sub>merge</sub> (%) | 11.5 (117.3) |
| R<sub>pim</sub> (%) | 3.2 (41.1) |
| CC<sub>half</sub> | 0.999 (0.784) |
| Mean I/signal | 19.1 (1.06) |
| Completeness (per cent) | 99.1 (86.6) |
| **Refinement Statistics** |  |
| Resolution (angstrom) | 29.95-1.88 |
| No. of reflections | 19678 |
| R<sub>work</sub>/R<sub>free</sub> (%) | 19.8/23.7 |
| Average B factor (Å<sup>2</sup>) | 33.71 |
| Wilson B factor (Å<sup>2</sup>) | 27.67 |
| **RMS deviation** |  |
| Bond lengths (Å) | 0.012 |
| Bond angles (°) | 1.044 |
| **Ramachandran Map** |  |
| Favoured (%) | 97.8 |
| Allowed (%) | 2.2 |
| Outliers (%) | 0 |

**Table 2.** Stoichiometry, affinity, and thermodynamic parameters corresponding to the ITC experiments. (averaged over n=3 repeats).

| | $K_d$ ($\mu M$) | $N$ | $\Delta H$<br>(kcal/mol) | $\Delta G$<br>(kcal/mol) | $-T\Delta S$<br>(kcal/mol) |
| --- | --- | --- | --- | --- | --- |
| MglA+RomX+GTP | $4.5 \pm 0.5$ | $1.1 \pm 0.1$ | $2.7 \pm 0.3$ | $-7.32 \pm 0.07$ | $9.9 \pm 0.3$ |
| RomR-REC<sup>125</sup>+RomX | $1.4 \pm 0.3$ | $1.3 \pm 0.2$ | $-1.6 \pm 0.3$ | $-8.1 \pm 0.02$ | $-5.31 \pm 0.3$ |
| RomR-REC<sup>141</sup>+RomX | $1.15 \pm 0.05$ | $1.3 \pm 0.2$ | $2.7 \pm 0.3$ | $-8.0 \pm 0.1$ | $-6.5 \pm 0.3$ |

### RomX fold is similar to the stalk domain of the dynamin family and is associated with response receivers

We used the experimentally determined structure of RomX (PDB ID: 8ZNA) to further identify proteins with a similar fold. Interestingly, the stalk domain of atlastin in humans and Sey1p in yeasts were found to be most structurally similar to RomX (Fig. 1C)^37–39^ based on output from DALI^40^ (example: RMSD 0.75 Å between 19 pruned atom pairs of PDB 4IDN, Supplementary Table S2). It is interesting to note that the occurrence of RomX fold in eukaryotes is associated with GTPases, considering that RomX (along with RomR) functions as a regulator of the prokaryotic small Ras-like GTPase MglA. However, the point of insertion between the stalk domain and the atlastin GTPase domain is away from the nucleotide-binding pocket, unlike interactions observed in typical GEFs, which sit near the nucleotide-binding pocket, competing out the bound GDP (Fig. 1C)^41–43^.

Further, using the experimental structure of RomX (PDB ID: 8ZNA), we performed a structure-based alignment using Foldseek^44^. Out of the hits from the AlphaFold database, many were models of uncharacterized bacterial proteins^45^ (Fig. 1D). Out of the annotated models, the most common hits were of response regulatory proteins containing a receiver domain (Fig. 1D, E) and a domain with RomX-like fold in tandem (RMSD 1.167 Å between 53 pruned atom pairs with AlphaFold model A0A2N6E6P0, a representative sequence from the list highlighted in green in Supplementary Table S2). This is an interesting observation in the context of *M. xanthus* RomX being associated with RomR which has an N-terminal response receiver (REC) domain. Another associated fold was of Zinc finger/ thioredoxin putative domain^46^ (Fig. 1D, E) which is associated with the REC-domain containing hits. This is interesting given the presence of a similar zinc finger domain (ZnR) in GltJ, a candidate protein within the *Myxococcus* focal adhesion complexes (FAC)^47^. The ZnR domain of GltJ has been shown to interact with MglB, significantly enhancing the GTP hydrolysis activity of MglA. The analysis reveals that motility-related complexes in *Myxococcus*, such as MglA/MglB, RomR/RomX, and FAC proteins, are prevalent across various deltaproteobacteria families, with a tendency for these domains to associate with one another.

### RomX forms a high-affinity complex with the RomR-REC domain

Next, we characterized the interface between RomX and RomR, and consequently attempted to dissect the precise domain boundaries of RomR that interact with RomX (Fig. 2A). For the N-terminal REC domain boundary, we chose residues 125 and 141 respectively as the domain boundarires for the two different constructs tested, based on secondary structure prediction (Supplementary Fig. S2A). We cloned and expressed the REC domain constructs (RomR-REC^125^ and RomR-REC^141^) of RomR (Fig. 2B) and recombinantly purified them (Supplementary Fig. S2B). A shorter construct of 1-116 residues was also attempted to be purified, based on the construct used in earlier reports^33^. However, it could not be successfully purified as the protein was unstable and probably aggregated during expression and purification steps.

**Figure 2.**
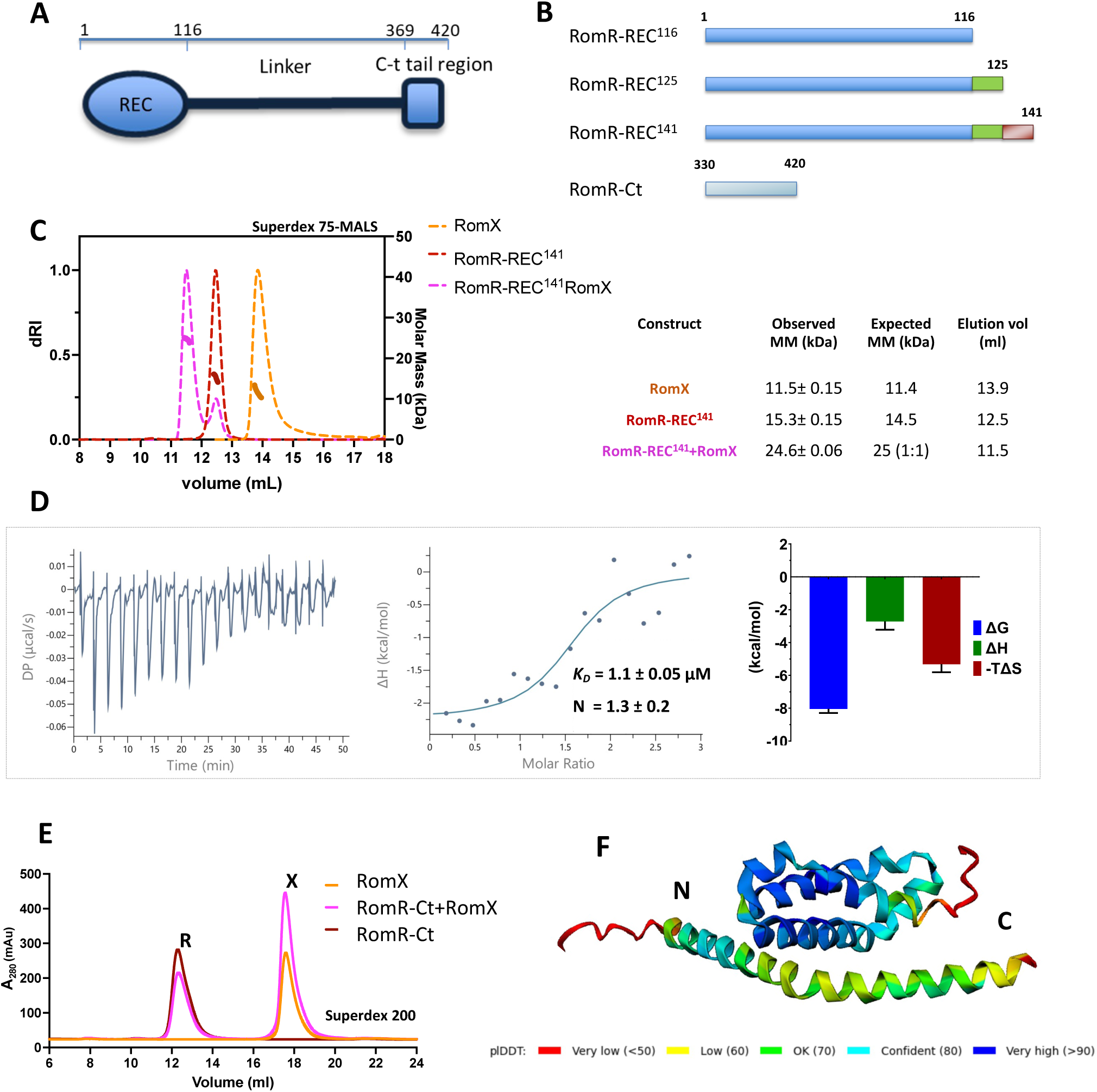
RomX interacts with RomR-REC and not with RomR-Ct. A: Schematic showing the domains of RomR, N-terminal REC domain, a long-disordered linker followed by a C-terminal helical domain. B: The domain-wise constructs of RomR that were designed for this study. The residue numbers of the respective boundaries are labelled. C: SEC run (using Superdex 75) coupled with multi-angle light scattering to show the interaction of RomR-REC^141^ and RomX. The molar mass of the corresponding peaks is plotted on the right y-axis. RomX is in orange, RomR-REC^141^ is in red, and complex is in magenta. The observed and expected molar mass and the respective elution volumes are tabulated on the right. D: ITC thermogram showing the titration of RomX with RomR-REC^125^. The left panel shows the raw data of exothermic heat pulses with time, the middle panel shows the corresponding differential binding curve fitted to a single site binding model and the right panel shows the thermodynamic signature plot detailing the ΔG, ΔH and -TΔS values corresponding to the same reaction (mean and standard errors are shown). E: Analytical SEC run using Superdex 200 showing no interaction of RomR-Ct with RomX. REC-Ct+RomX (orange), RomR-Ct (brick red), RomR + RomX (magenta). F: AlphaFold model of RomR-Ct in complex with RomX. The low pLDDT scores are highlighted in the regions labeled in green and yellow.

The purified constructs of RomR-REC domain were subjected to analytical size exclusion chromatography with RomX. The RomR-REC^125^ in complex with RomX, resulted in a shift in the resultant peak to a higher volume (12.1 ml), as compared to individual peaks of RomR-REC^125^ (12.9 ml) and RomX (13.8 ml). This confirms the complex formation (Supplementary Fig. S3A). To determine the stoichiometry of the complex, we performed SEC-MALS to determine the molar mass corresponding to the peak of the complex (Fig. 2C). Using a Superdex 75 SEC column, there was a shift in the peak of the complex from 12.5 ml for RomR REC^141^ and 14 ml for RomX to 11.5 ml. The observed molar mass was 24.6 kDa, corresponding to a 1:1 complex of RomR REC^141^ (14.5 kDa) and RomX (11.4 kDa).

Further, the binding affinity between RomR-REC^125^ and RomX complex was quantified using isothermal titration calorimetry (Fig. 2D). A similar titration experiment was performed with RomR-REC^141^, and the results were comparable. We obtained an average *K_D_* of 1.14 ± 0.05 μM for RomR-REC^125^ and RomX and 1.2 ± 0.5 μM for RomR-REC^141^ and RomX (Fig. 2D). The stoichiometry of interaction was the same for both RomR-REC^125^ and RomR-REC^141^, 1.3 ± 0.2 and 1.2 ± 0.2, respectively, implying an interaction of one molecule of RomR-REC domain with one molecule of RomX (Supplementary Fig. S2C). The interaction was observed to be exothermic, and entropy-driven. Based on our comparable results between RomR-REC^125^ and RomR-REC^141^, we can conclude that residues 1-125 consist of the minimal functional REC domain in RomR, capable of maintaining a stable RomX interaction.

Next, the C-terminal domain, spanning residues 331-420 (based on secondary structure prediction), was cloned and successfully purified (Supplementary Fig. S2D). The C-terminal domain was found to be predominantly composed of alpha-helices as confirmed by CD spectroscopy (Supplementary Fig. S2E). According to secondary structure and AlphaFold based predictions, RomR-Ct helical domain comprises helices from residues 371-420, which is kinked in the center around the 396^th^ residue (Fig. 2F). Based on analytical SEC (Superdex 200), it was observed that the RomR-Ct helical domain does not interact with RomX. There was a clear separation of peaks observed at 12.3 ml for RomR-Ct and 17.6 ml for RomX (Fig. 2E, Supplementary Fig. S2F). Further, using AlphaFold, we could not predict a reliable interface between RomR-Ct domain and RomX due to low pLDDT scores (Fig. 2F, Supplementary Fig. S4)^37,48^. The SEC result indicates that the C-terminal helical domain alone of RomR does not form a stable complex with RomX.

### Salt bridge interactions predominantly hold RomX and RomR-REC complex

To characterize the interface between the RomR-REC domain and RomX, we attempted to crystallize the complex of RomX with RomR-REC^125^. We majorly obtained crystals of RomX and phase-separated droplets of RomR in the crystallization drops. Hence we proceeded to predict an AlphaFold model of the complex using one molecule each of RomX and RomR-REC^125^ (Fig. 3A, Supplementary Fig. S4). The predicted RomX model superposed well with the experimentally obtained RomX structure (PDB ID: 8ZNA) with an RMSD of 0.8 Å. Analysis of the interface from the AlphaFold prediction of the complex, Glu-108 of RomR formed a salt bridge interaction with Arg-76 of RomX. Further, Val-17 of RomR and Leu-75 of RomX formed a hydrophobic core into which Trp-87 of RomX stacks when in complex with RomR-REC. This tryptophan is exposed in solution in the crystal structure when RomX is monomeric. We checked the extent of conservation of these candidate interface residues both in RomR-REC and RomX. The Glu-108 in RomR is highly conserved (Fig. 3B), and so is the Trp-87 in RomX (Fig. 3C). The Arg-76 of RomX is often replaced with Lys, which highlights the importance of these basic residues at this position capable of forming the RomR-REC/RomX interface. It is important to note that according to the model, the candidate interface residues in RomX are part of helix-3.

**Figure 3.**
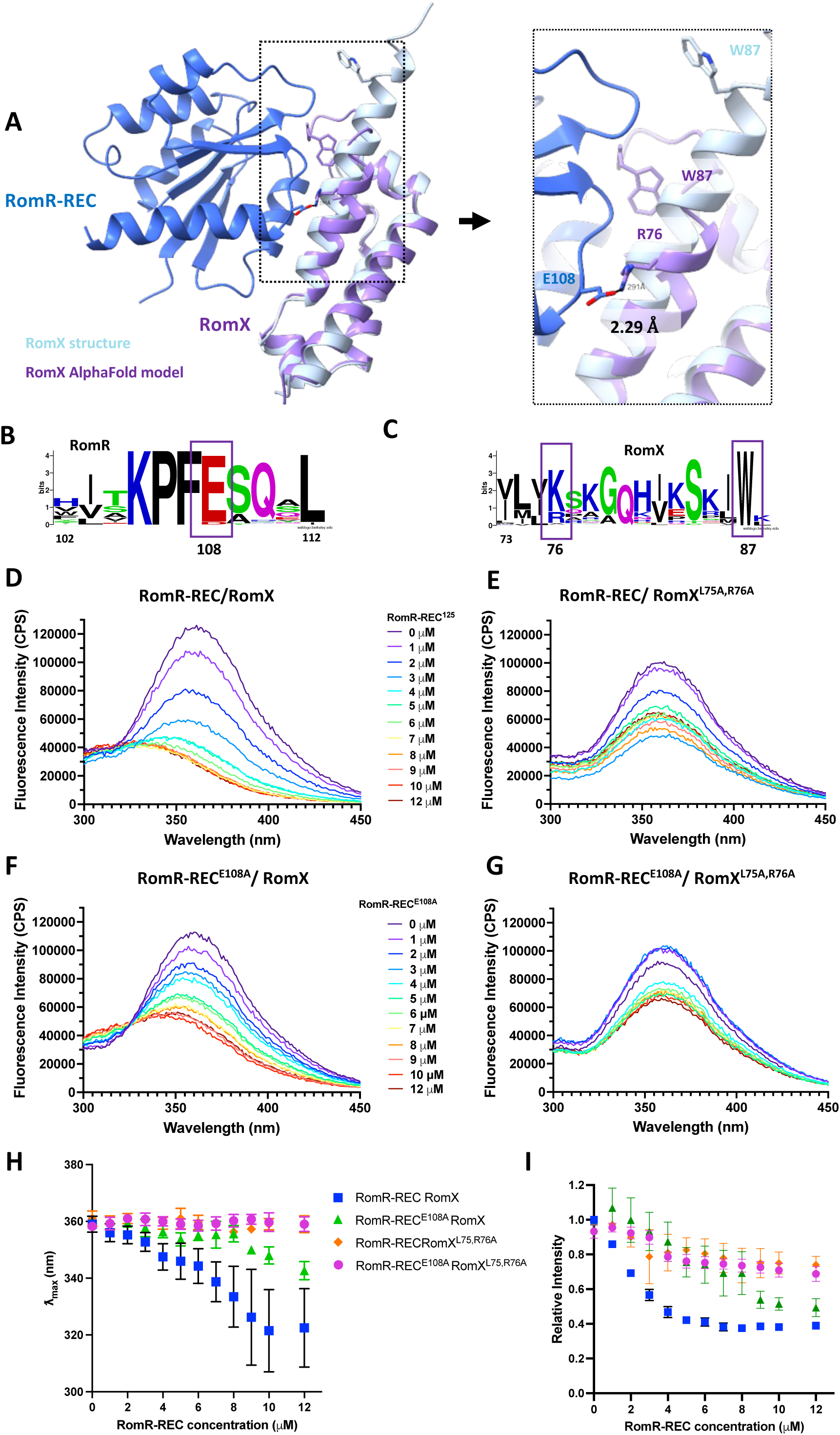
Interface residues between RomR-REC and RomX are held by salt-bridge interaction. A: AlphaFold model of RomR-REC (blue) and RomX complex. Experimentally obtained RomX structure in blue-grey is superposed on the model in purple. On the right, the interface of the complex is expanded showing the probable interface residues, W87 and R76 in RomX and E108 in RomR. The interacting distance between RomR E108 and RomX R76 is highlighted. The conformational change of RomX W87 across the monomeric (blue) and complex states (grey) is also shown. B, C: WebLogos showing the conservation of residues in RomR-REC and RomX interfaces, respectively. The candidate residues are highlighted and their positions are labeled below. D, E: Tryptophan fluorescence spectra to determine the complex formation WT RomR-REC and RomX in WT and RomX^L75A,R76A^ (left and right, respectively). RomX is titrated with increasing concentrations of RomR-REC as shown in the legend. F, G: Tryptophan fluorescence spectra to determine the complex formation RomR-REC^E108A^ mutant and RomX in WT and RomX^L75A,R76A^ (left and right, respectively). RomX is titrated with increasing concentrations of RomR-REC^E108A^ as shown in the legend. H, I: Relative fluorescence intensity at emission maxima Lmax and the corresponding Lmax values are plotted against the concentration of RomR-REC to determine the shift in fluorescence spectra as well as quenching due to complex formation, due to change in the environment of the Trp87 at the RomRX interface. The mean of 3 replicates is plotted with the error bars indicating standard errors.

Since Trp-87 was found to be the sole tryptophan residue in the RomRX complex showing a conformational transition across the monomeric RomX and RomRX complex, it could be used as a probe to monitor RomRX complex formation. The structure prediction was validated by monitoring the fluorescence of the single Trp-87 in RomX. Trp-87 is probably exposed in monomeric RomX and is buried in the RomRX complex interface. This would show up as a shift in the wavelength and intensity of the Trp emission maximum. Hence, RomX was titrated with increasing concentrations of RomR-REC^125^ (Fig. 3D). There was indeed a decrease in the Trp fluorescence intensity and a gradual blue shift in the emission maximum from 360 nm to near 320 nm (Fig. 3H, I). This indicated a reduction in the solvent-exposed environment of the Trp as the RomRX complex formed. This confirms that Trp-87 is one of the residues that inserts into the RomRX complex interface, a feature consistent with the AlphaFold prediction.

Based on our understanding of the interaction interface, we designed specific mutants of RomR and RomX, namely, RomR-REC^E108A^ and RomX^L75A, R76A^ (where both L75 and R76 were mutated to alanines). Since RomR-REC^125^ was identified as the minimal functional domain interacting with RomX, the E108A mutation was introduced in the RomR-REC^125^ background (Supplementary Fig. S5A). We then measured tryptophan fluorescence using the interface mutants of RomR-REC^125^ and RomX. The RomX^L75A, R76A^ mutant was titrated with increasing concentrations of wild-type RomR-REC^125^ (Fig. 3E). No spectral shift was observed compared to the wild-type RomR-REC^125^ and RomX interaction (Fig. 3H). However, there was a slight reduction in the intensity of the emission maxima at higher concentrations of RomR-REC^125^, likely due to molecular crowding, which could partially mask the Trp-87 fluorescence signal (Fig. 3I). This suggests that RomX Arg-76, which is oriented toward RomR Glu-108, plays a critical role in complex formation, thereby stabilizing the interface where Trp-87 stacks.

When wild-type RomX was titrated with increasing concentrations of mutant RomR-REC^E108A^, a decrease in Trp fluorescence intensity and a slight blue shift in the emission maximum to approximately 340 nm were observed (Fig. 3F, H, I). However, this effect was less pronounced compared to the interaction between wild-type RomR-REC^125^ and RomX. Furthermore, when we titrated the RomX^L75A, R76A^ mutant with the RomR-REC^E108A^ mutant, there was no spectral shift in the emission maxima, and the fluorescence quenching observed was similar to that of the RomX^L75A, R76A^ mutant alone.

To further validate this, we performed ITC measurements to quantify the binding between RomR-REC^E108A^ and RomX^L75A, R76A^ (Supplementary Fig. S5B). No binding was observed between these mutant proteins. Together, these results highlight the role of RomX in facilitating proper complex formation, with the Arg-76 residue stabilizing the RomR-RomX interaction through a salt bridge with RomR Glu-108.

### RomX forms a complex with MglA only in the presence of GTP

Next, we attempted to decipher the preferred nucleotide state for the interaction of MglA with RomX. Analytical SEC was carried out using MglA and RomX in the presence of GDP and GTP, respectively (Supplementary Fig. S6A, B). MglA and RomX eluted at 13.6 ml and 14 ml, respectively. These elution volumes of the peaks were too close to be resolved conclusively. Since there was no shift of the resultant peak of a complex of MglA and RomX towards a lower elution volume, we could not demonstrate any interaction between the two proteins in the presence of either nucleotide. Considering the plausible effects of dilution and aberrant elution volumes of the complex, we could not conclusively determine if RomX binds to MglA either in the presence of GTP or GDP.

To resolve this, we performed isothermal titration calorimetry (ITC) studies to quantify the interaction between MglA and RomX in the presence of nucleotides. There was no binding observed between MglA and RomX in the presence of GDP unlike what would be expected for a typical GEF (Supplementary Fig. S6C). Instead, when we performed titration in the presence of excess GTP (2 mM), we observed binding between MglA and RomX (Fig. 4A). We observed a *K_d_* of 4.8 ± 0.5 μM and the stoichiometry of interaction N was 1.1 ± 0.1. This confirmed that RomX interacts with MglA in a 1:1 ratio in the presence of GTP. Based on the observed ΔH and ΔS values, the interaction is entropy-driven (Fig. 4A). The positive ΔH also rules out the possibility that the observed energy changes are the result of GTP hydrolysis, which is negligible in the absence of MglB.

**Figure 4.**
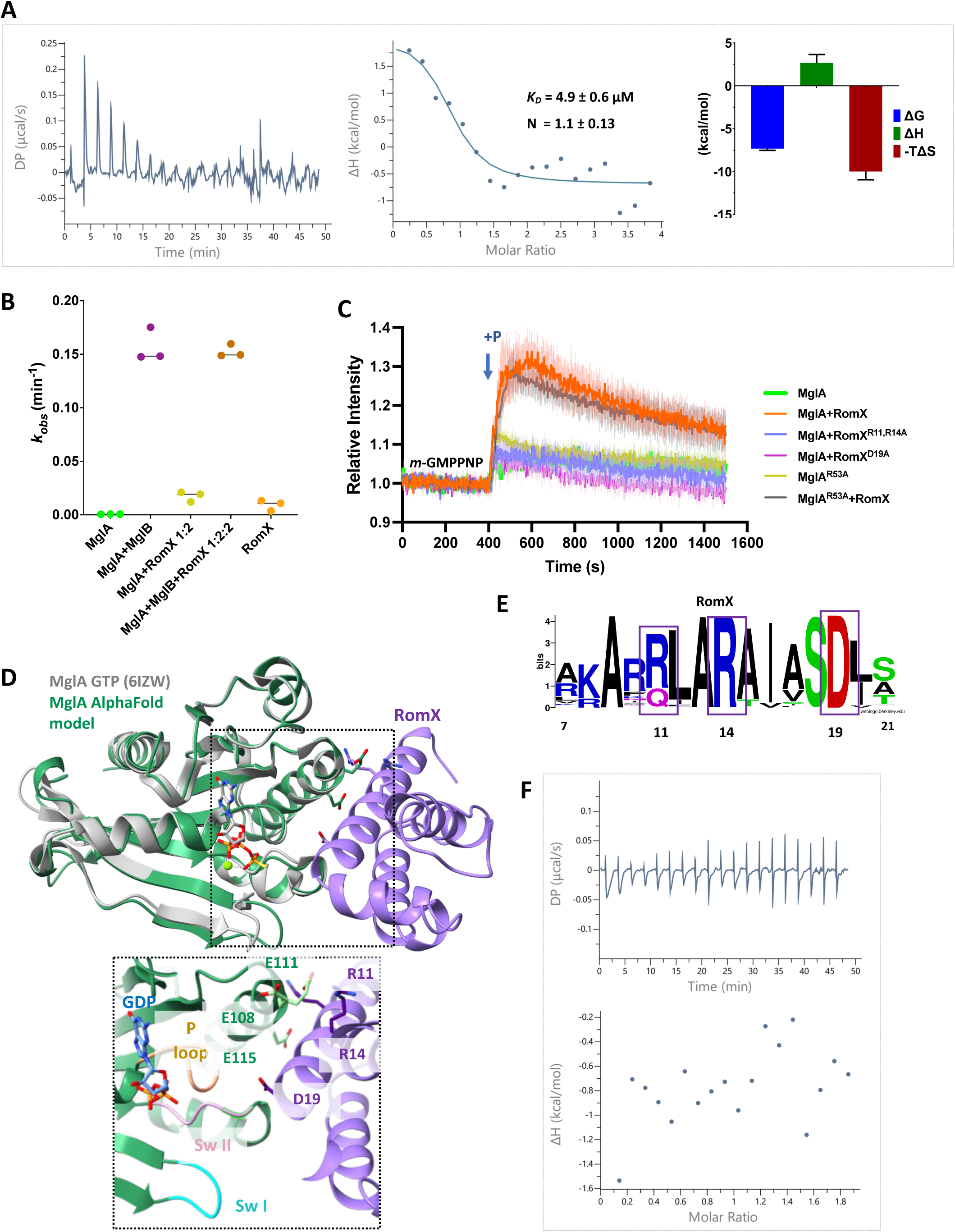
RomX preferentially interacts with MglA-GTP near the nucleotide-binding pocket, stimulating exchange. A: ITC thermogram of a representative binding assay showing the titration of RomX with MglA in the presence of GTP and magnesium. The left panel shows the raw data of endothermic heat pulses with time. The middle panel shows the corresponding differential binding curve fitted to a single-site binding model. The right panel shows the thermodynamic signature plot detailing the ΔG, ΔH, and -TΔS values corresponding to the same reaction (mean and standard errors are shown). B: NADH coupled GTPase assay data showing the k_obs_ values corresponding to GDP turnover by MglA in the presence of RomX. MglA with MglB is shown as a positive control. The black line represents the mean and the coloured bars represent the standard error. C: Plots (from n=3 repeats) showing *mant*-GMPPNP association assay of MglA with RomX, where the darker lines represent the mean of three replicates and the lighter traces represent the respective standard errors. At 400 s, the protein was added, as shown by the arrow. RomX (in orange) stimulates the release of GDP on MglA and uptake of *mant*-GMPPNP in a 1:2 ratio (MglA 3 μM: RomX 6 μM), whereas MglA alone (in green) does not show exchange. Similarly, RomX stimulates the release of GDP on MglA^R53A^ mutant and uptake of *mant*-GMPPNP in a 1:2 ratio (in grey), whereas MglA^R53A^ alone (in ochre) do not show exchange The RomX mutants at the MglA interface (RomX^R11,14A^ in magenta and RomX^D19A^ in blue) also show abrogated nucleotide exchange activity. D: AlphaFold model of MglA (in green) with RomX (purple) with MglA-GTP (in gray) superimposed. In the bottom inset, the interface residues are highlighted (MglA residues in green and RomX residues in purple). P-loop, Switch I, and Switch II of MglA are highlighted in orange, pink, and cyan, respectively. E: WebLogo showing the conservation of residues in RomX interfacing with MglA. The candidate residues are highlighted and their positions are labeled below. F: ITC thermogram of a representative binding assay showing the titration of RomX^R11,14A^ with MglA in the presence of GTP and magnesium. The top panel shows the raw data of heat pulses with time. The lower panel shows the corresponding heat exchange, which could not be reliably fitted to a binding curve, indicating no binding

Since we observed the binding of RomX with MglA in the presence of GTP, we estimated the effect of RomX on MglA GTP hydrolyzing activity to rule out the possibility of a GTPase-activating protein (GAP) activity. There was no stimulation in the MglA GTP hydrolysis upon the addition of RomX (Fig. 4B). The results were similar even when higher concentrations of RomX were used (1:4 MglA: RomX). MglA+MglB (1:2) was used as a positive control where the GAP stimulation of MglA was observed, with no further stimulation with RomX.

### RomX helix-1 interacts with MglA-GTP, stimulating nucleotide exchange

We proceeded with the measurement of the nucleotide exchange activity of the purified RomX as reported earlier^32^. Nucleotide exchange assays were performed where we monitored the *mant*-GMPPNP association (Fig. 4C). It was consistently observed that in the presence of RomX, MglA-GDP (purified MglA comes bound to GDP^21^) showed robust nucleotide exchange and an increase in fluorescence was observed upon addition of MglA with RomX to *mant*-GMPPNP, compared to the addition of MglA alone. This demonstrated that RomX exerts a nucleotide exchange activity on MglA, in line with earlier reports. However, the interaction is probably not stable as we observe an intrinsic decrease in fluorescence intensity immediately after the association event. This also confirms that RomX indeed interacts with MglA, consistent with the ITC measurements, however, the interaction is not sufficient to exert a stabilization effect of the bound nucleotide conformation of MglA. This is suggestive that the interaction might be transient given the comparatively high *K_d_*.

Based on the interaction studies between MglA and RomX, we wanted to characterize the interface between MglA and RomX. Crystallization trials were unsuccessful to date possibly because of the transient nature of the complex. Hence, we used AlphaFold to generate a complex of MglA with RomX (Fig. 4D, Supplementary Fig. S4)^49^. The helix-1 from the N-terminal of RomX was seen to face toward the nucleotide-binding pocket of MglA. There are arginines (Arg-11, Arg-14) of RomX and glutamates (Glu-108, Glu-111, Glu-115) of MglA, which come in close proximity and could be holding the complex together. Moreover, Asp-19 faces towards the MglA nucleotide binding pocket near the Switch II, which could be the possible residue driving the nucleotide specificity and exchange activity on MglA. Interestingly, all the 3 chosen interface residues, Arg-11, Arg-14, and Asp-19 of RomX are conserved (Fig. 2E), which indicates that these might be important for driving RomX function.

Arg-53 is the Switch 1 residue of MglA, which is known to stabilize the GTP-bound conformation of MglA^17,20,21^. To determine whether the interaction of RomX with MglA-GTP is specifically mediated through Switch 1 Arg-53, we examined the nucleotide exchange activity of the MglA^R53A^ mutant (Fig. 4C). This mutant had been previously studied in the context of MglB interaction and here, we checked its role in the interaction with RomX. The increase in fluorescence observed with the MglA^R53A^ mutant in the presence of RomX was comparable to that seen with wild-type MglA. As with wild-type MglA, MglA^R53A^ alone did not exhibit nucleotide exchange activity. This suggests that the specific interaction between RomX and MglA-GTP does not involve the Switch 1 Arg-53 residue.

To confirm the RomX-MglA interface and the consequent mechanism of RomX GEF activity, we generated mutants of RomX. Firstly, we generated a double mutant where the Arg-11 and Arg-14 of RomX were mutated to alanines, intending to break the transient interaction with MglA (RomX^R11,R14A^). We further mutated the Asp-19 of RomX, to alanine to abolish the GEF activity (RomX^D19A^) (Fig. 4D). We purified the mutant proteins (Supplementary Fig. S6D) and confirmed using CD spectroscopy that they were well folded (Supplementary Fig. S6E). We then checked their respective GEF activity in the presence of MglA using nucleotide exchange assays (Fig. 4C). It was observed with our *mant*-GMPPNP exchange assays that, indeed, both the RomX mutants had their GEF activity completely abolished as compared to the wild-type RomX. We additionally performed ITC experiments for monitoring the binding of RomX^R11,R14A^ with MglA in the presence of GTP (Fig. 4F). We observed no binding of this mutant to MglA-GTP, confirming that these residues in RomX were indeed involved in MglA interaction. For the RomX^D19A^ mutant, the overall protein yield after purification was significantly lower, although the corresponding CD spectra confirmed that the protein was properly folded. Due to the weak affinity between MglA and RomX, higher stock concentrations of RomX (1 mM) required for the ITC experiments could not be achieved for RomX^D19A^ using the experimental protocol for ITC. Hence, we were unable to reliably determine whether RomX^D19A^ can bind to MglA using ITC, however, the loss of GEF activity is suggestive of Asp-19 being at the putative interface between MglA and RomX.

### RomR-REC does not enhance the nucleotide exchange activity of RomX

We performed analytical SEC to dissect the interaction of the RomR-REC/RomX complex with MglA. No coelution was observed when the RomR-REC^125^/RomX complex was injected with MglA. There was elution at approximately 12.1 ml corresponding to the RomR-REC^125^/RomX complex and at 13.1 ml to MglA, as verified by SDS PAGE to resolve the overlapping peaks (Supplementary Fig. S3A). This observation was consistent when the same experiment was performed in the presence of GDP and GTP, respectively. A similar result was even observed when the same set of runs were performed with RomR-REC^141^. This observation is consistent also with the SEC profile for monitoring MglA-RomX interaction.

Further, we investigated the interaction of RomR-Ct with MglA in analytical SEC (Supplementary Fig. S3B). There was a clear separation of peaks of RomR-Ct (12.8 ml) and MglA (around 16 ml), both in the presence or absence of nucleotides. This shows that there is either no interaction or only a weak interaction of RomR-Ct with either RomX or MglA. All these observations indicate that there is probably no direct interaction between MglA and the RomR-REC domain or the C-terminal tail.

Based on our information regarding the interface between MglA-RomX and RomX-RomR-REC, we conclude that RomX is probably sandwiched between MglA and the RomR-REC domain on either side. We used AlphaFold to confirm our model of MglA : RomX : RomR-REC complex. The AlphaFold model supports our idea that RomR-REC scaffolds RomX which further recognises and interacts with MglA-GTP (Fig. 5A, Supplementary Fig. S4).

**Figure 5.**
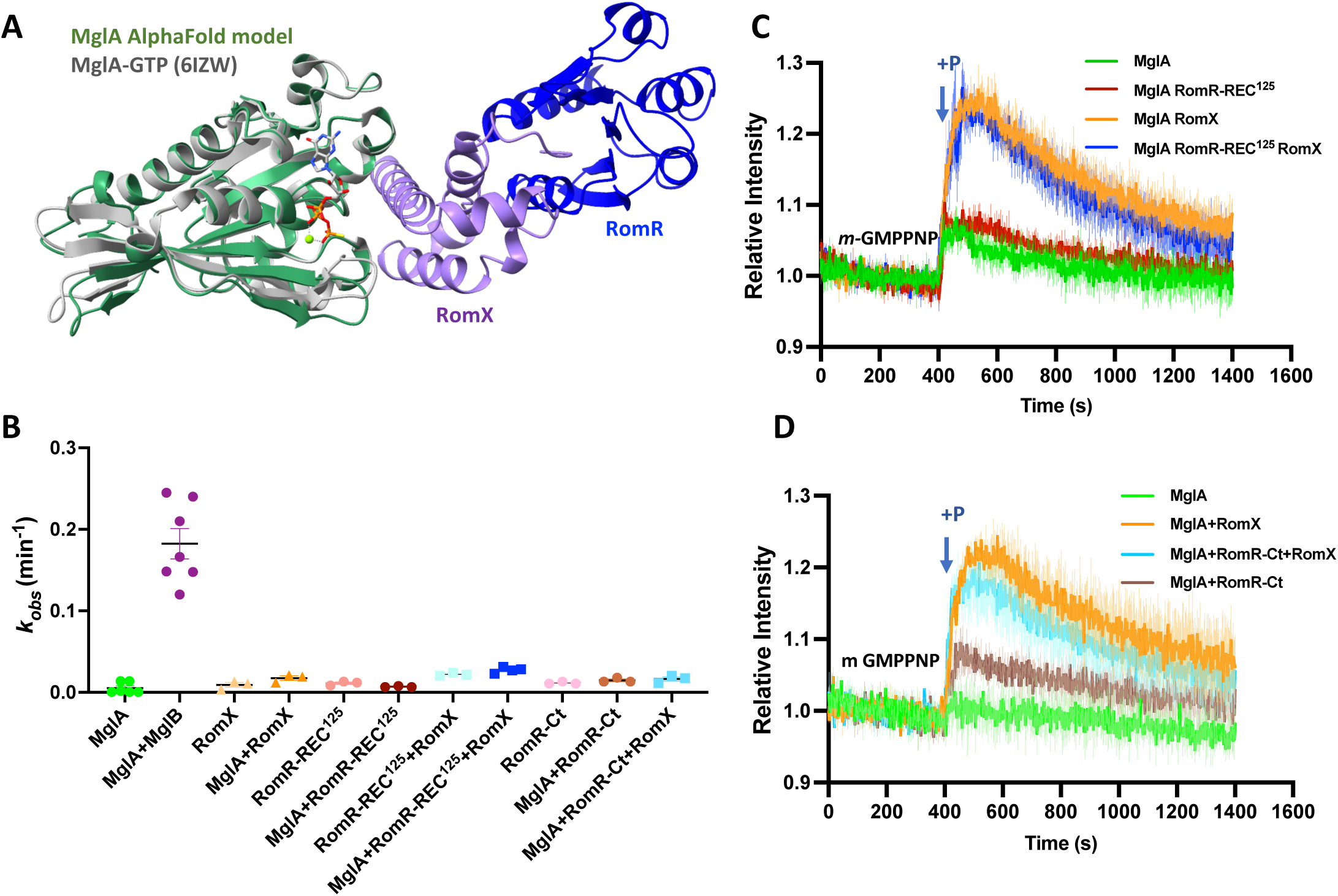
RomX is sandwiched between MglA and RomR-REC. A: AlphaFold model of MglA (in green), RomR (in dark blue) and RomX (in purple) predicts RomX sandwiched between MglA and RomX. MglA model is superposed with MglA-GTP (PDB ID: 6IZW, in gray). B: NADH coupled GTPase assay data showing that none of the RomR constructs with or without RomX shows GAP activity as observed with MglB. The black line represents the mean and the coloured bars represent the standard error. C and D: Plots showing *mant*-GMPPNP association assays of MglA with RomR REC^125^, and RomR-Ct domains, respectively, where the darker lines represent the mean of three replicates and the lighter traces represent the respective standard errors. The trend was similar for both cases. RomX (orange) stimulates the release of GDP on MglA and uptake of *mant*-GMPPNP in a 1:2 ratio and the rates are similar with RomX+ RomR (blue) in the same 1:2 ratio (MglA 3 μM : RomRX 6 μM), whereas RomR alone (brick red) does not stimulate exchange. n=3 repeats performed for each MglA, MglA+RomX, MglA+RomR-REC^125^, MglA+RomR-REC^125^+RomX, MglA+RomR-Ct, MglA+RomR-Ct+RomX, runs.

We proceeded to check the role of all the RomR constructs in MglA GTPase activation (Fig. 5B) with and without RomX. It was observed that RomR-REC domain constructs and RomX alone or as a complex did not accelerate the rates of GTP hydrolysis by MglA. Expectedly, RomR-Ct also did not stimulate the GTPase activity of MglA. This indeed confirms that there is no stimulation of MglA-GTP hydrolysis by the RomR-RomX complex.

Since we observed the interaction of RomR-REC with RomX, we wanted to check if the presence of RomR-REC stimulates the GEF activity of RomX. We monitored the exchange of GDP bound to MglA in the presence of *mant*-GMPPNP (Fig. 5C). RomX induced a spontaneous nucleotide exchange on MglA, at similar rates with and without RomR-REC^125^.

RomR-REC^125^ alone did not stimulate nucleotide exchange by MglA. This showed that RomX alone drives nucleotide exchange on MglA without any additional contribution from the RomR REC. Expectedly, we did not observe any effect of RomR-Ct in the GEF stimulation of MglA (Fig. 5D). Hence, in conclusion, RomX forms a complex with RomR-REC and MglA in the presence of GTP, confirming that RomX is sandwiched between its two interactors, with no direct interaction between MglA and RomR-REC. This is in line with the previous hypothesis based on *in vivo* studies^32^.

## Discussion

We report the experimentally derived structure of RomX and identified the fold to be composed of a three-helix bundle (PDB ID: 8ZNA). Our analyses revealed that the RomX fold is most often associated with a response receiver domain, which is prevalent in bacteria. Many of these proteins remain uncharacterized functionally. This observation suggests that the interaction between the RomX-like fold and the receiver domain is conserved and likely integrated as a part of these individual response regulatory proteins. In the future, investigating deeper into the mechanism of function of these multidomain response regulators and the role of the three-helix bundle RomX-like fold in these proteins will be of great interest.

We further identified that the helical stalk domain of the dynamin family of GTPases is a structural homolog of RomX. The stalk domain in atlastin, especially, is shorter and closer to RomX owing to the presence of a three-helix bundle^37,48^. This domain is involved in the dimerization of the GTPase exclusively in the GTP-bound state to trigger hydrolysis driving membrane fusion^50^. The conformational changes in the stalk domain with respect to the GTPase domain in atlastin across nucleotide-bound states offer an interesting analogy in light of the specificity of RomX to MglA exclusively in the GTP-bound state. However, the atlastin stalk domain triggers nucleotide specificity for dimerization by altering the conformation of the switch regions through intramolecular interactions^32^. In contrast, RomX likely interacts near the nucleotide-binding pocket of MglA via helix-1, according to the AlphaFold model, and further validated by our mutational studies in RomX. These residues also appear to be highly conserved, suggestive of a functional role for them. Confirmation of this interaction can be achieved through complementary mutations in MglA near the interface where RomX helix-1 interacts. However, mutating residues in MglA near the nucleotide-binding pocket often significantly destabilizes the protein, making binding studies such as ITC challenging to perform and interpret. We checked the role of RomX in stabilizing the GTP-bound conformation of MglA through the Switch 1 motif by using the previously characterized MglA^R53A^ mutant. However, our results indicate that RomX does not directly interact with the Switch 1 motif, consistent with the AlphaFold model, which shows RomX positioned far from the switch residues.

In line with the previous report, we observe that wild-type RomX stimulates nucleotide exchange activity on MglA. Additionally, the RomX helix-1 mutants abolish the exchange activity of RomX, further confirming the significance of this interface in nucleotide specificity and interaction with MglA. However, we do not detect any interaction between RomX and MglA-GDP, as would be expected for a typical GEF. This is again consistent with the previous report^32^. Based on these findings, we conclude that RomX predominantly interacts with and sequesters GTP-bound MglA thereby transiently increasing the population of GTP-bound MglA (Fig. 6A). However, currently, the structural basis of nucleotide specificity of RomX to MglA-GTP still needs to be investigated further. Owing to the transient nature of the complex, interaction with other stabilizers could assist structural studies in the future which could answer this question.

**Figure 6.**
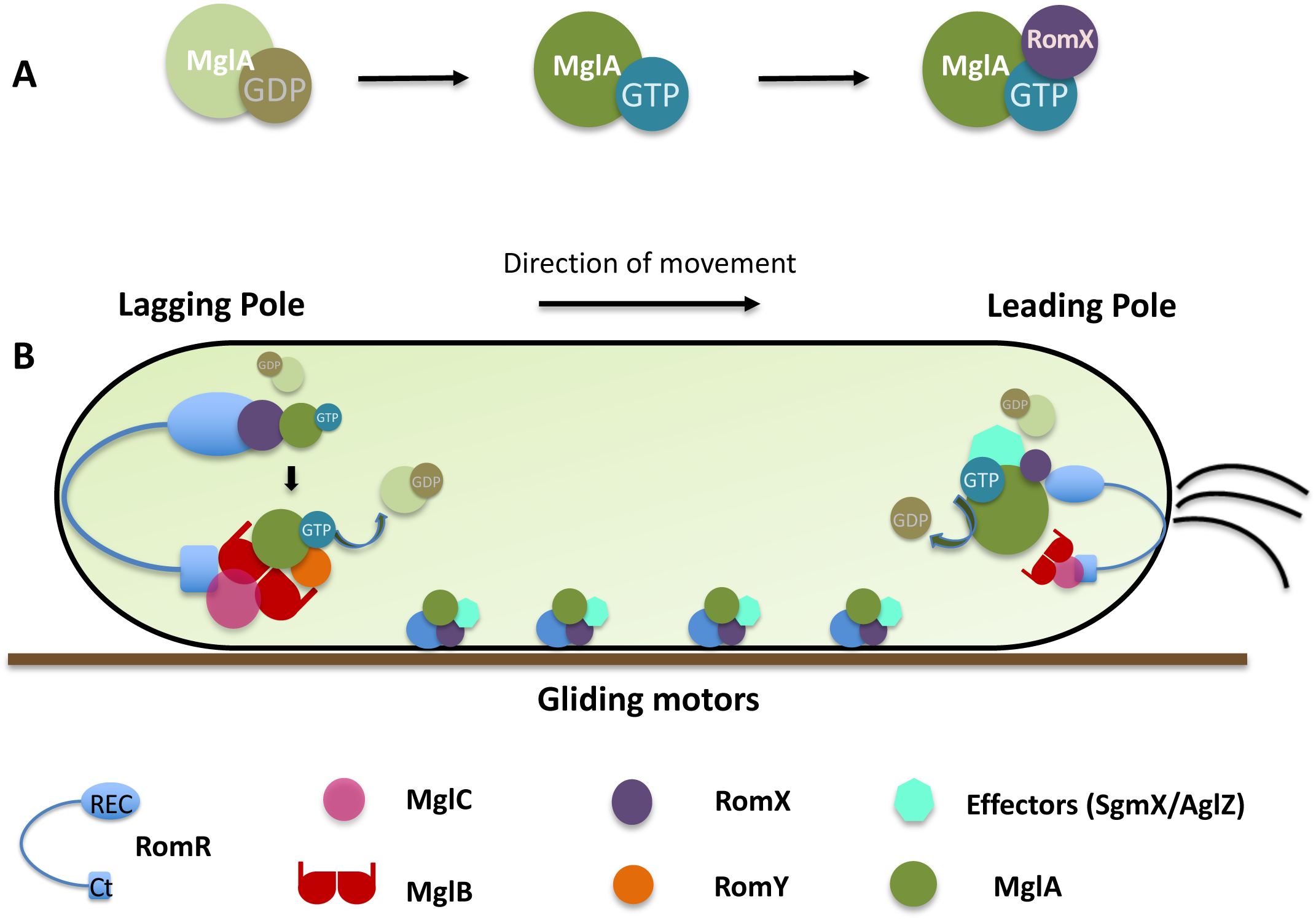
RomX maintains *M. xanthus* cell polarity by regulating MglA localization. A: Mechanism of stimulation of nucleotide exchange on MglA by RomX. RomX preferentially binds and sequesters MglA-GTP facilitating nucleotide exchange. B: Model showing the mechanism of polarity maintenance in *M. xanthus*. At the lagging pole RomX (in purple) generates local pools of MglA-GTP, ready to be hydrolyzed by MglB-RomY, thereby excluding MglA. At the leading pole, RomX recruits MglA-GTP which is probably facilitated by interaction with additional effectors (in teal).

Canonical GEFs for small Ras-like GTPases typically interact with the GTPase specifically in its GDP-bound conformation. On the other hand, specificity to the GTP-bound state is frequently exhibited by effectors, thereby facilitating downstream cellular functions. For instance, the structure of PDZ-RhoGEF (PRG), a member of the RGS-RhoGEFs, in complex with GTP-bound RhoA reveals an interface strikingly similar to the canonical GTPase-effector interface^51^. RalGDS, initially believed to act as a GEF for the small GTPase Ral by promoting the exchange of GDP for GTP and thus activating it, was also found to function predominantly as an effector of the small GTPase Ras^52,53^. Furthermore, while most characterized GEFs activate their GTPases by closely interacting with the switch residues of the GTPase, preliminary results suggest that RomX’s interaction with MglA does not rely on these switch residues^42^. This observation raises the possibility that RomX may unconventionally function as an effector or stabilizer of GTP-bound MglA rather than as a traditional GEF, or may even have a dual role.

Our binding experiments demonstrate that helix-3 of RomX interacts with the RomR-REC domain, facilitated by salt-bridge interactions involving conserved residues in RomX helix-3 and RomR-REC. Given that RomX interacts with MglA via helix-1, positioned opposite to helix-3, it supports the idea that RomX is likely sandwiched between MglA and RomR-REC. Notably, we did not detect a direct interaction between MglA and RomR-REC. This observation is validated by the lack of further acceleration in the nucleotide exchange activity of RomX in the presence of RomR-REC. Conducting further *in vivo* experiments to assess the effects of interface mutants, as predicted in this study, will provide insights into the mechanism through which the RomR/X module contributes to maintaining MglA localization in a polarised cell.

We have determined that residues 1-125 constitute the minimal folded functional REC domain in RomR. Furthermore, we conclude that the REC domain of RomR interacts with RomX in a 1:1 ratio. This explains why an exact co-localization of RomR with RomX in a bipolar asymmetric fashion is observed as per *in vivo* studies^32^. We hypothesize that the localization of RomX is largely a consequence of the asymmetric bipolar localization of RomR. Specifically, the REC domain of RomR recruits RomX, in a 1:1 ratio to the cell poles, while the C-terminal output domain aids in recruiting other polar candidates^54^. Hence RomX localization simply mirrors the RomR localization pattern.

In the cellular context, a pertinent question arises: during polarity maintenance, why does a bigger RomR/X cluster persist at the lagging cell pole when its function as an MglA-GTP recruitment factor is primarily exerted at the leading pole? It is to be noted that RomR serves as a polar scaffold not only for RomX but also for other polarly localized proteins such as MglC and MglB^54–57^, thereby regulating their dynamics and function. The substantial cluster of RomR at the lagging pole plays a crucial role in maintaining a refractory period due to its slower dynamics. This ensures that a cell cannot be immediately triggered again after reversal^34^. We presume that there could also be a function of RomX localized at the lagging pole. MglA-GTP exhibits a higher affinity for MglB compared to RomX (approximately 5-fold), and this difference may be further accentuated in the presence of RomY exclusively at the lagging cell pole^21,58^. This GAP complex actively triggers GTP hydrolysis, excluding MglA-GDP from the lagging pole in between reversals. Any excess MglA-GTP is transiently stabilized by RomX until the MglB/RomY complex becomes available to hydrolyze MglA, collectively facilitating the active exclusion of MglA from the lagging pole (Fig. 6B).

The function of RomX as a MglA-GTP recruiter is primarily manifested at the leading pole and the Agl/Glt complexes, as previously hypothesized^32^. However, as this complex is inherently unstable, as per our observations, additional effectors such as SgmX, AglZ, or PilB may be required to stabilize the recruitment of MglA-GTP near the leading pole^22,23,28,59^ (Fig. 6B). During polarity switching, the RomX population at the old lagging pole could actively recruit MglA-GTP in the absence of MglB, with the help of effectors. This recruitment could help form a new cluster of MglA-GTP, defining the new leading pole.

We also identified a zinc finger (ZnR) domain-containing fold from our analyses, which is predominantly associated with response receiver domains. GltJ, a FAC candidate, bears a ZnR domain that has been shown to interact with MglB and trigger MglA GTP hydrolysis at the lagging cell pole^47^. This is critical in disassembling the FACs at the lagging cell pole. Together, the structural characterization of RomX has helped us identify proteins containing domains of related folds among both eukaryotes and prokaryotes suggesting a prevalent functional association. It is interesting to observe the existence of RomX fold associated with the response receiver domains in prokaryotes and to GTPases in higher eukaryotes, an observation that may hold significance in the evolution of P-loop NTPases and cellular signalling mechanisms.

## Materials and Methods

### Cloning

Restriction-free cloning was performed for all the constructs. For the REC domain of RomR, residues 1-115 of the RomR gene was amplified from *romR-pETPhos* (obtained from Dr. Tam Mignot’s lab) construct. It was cloned in *pHis17-Kan^R^* vector with a C-terminal hexahistidine tag using respective primers (Supplementary Table S3). Further, a construct with N-terminal hexahistidine tag was cloned. Since the protein was impure, further 1-125 and 1-141 residues of the *romR* gene were cloned in the *pHis17-Kan^R^*vector with a C-terminal hexahistidine tag. For the C-terminal, residues 331-420 *romR* gene were cloned in the *pHis17-Kan^R^* vector with a C-terminal hexahistidine tag (insert amplified from *romR* gene synthesized from BioTechDesk). RomX gene (MXAN_3350) was amplified from *M. xanthus* genomic DNA (obtained from Dr. Tam Mignot’s lab) and cloned into *pHis17-Amp^R^* vector with both N and C-terminal hexahistidine tags. A similar strategy as mentioned above was also used to generate interface mutants of RomR and RomX. For the RomR-REC mutants, a RomR-REC^125^ background was used. MglA and MglB constructs used were as reported in Baranwal et al, 2019^21^. All clones were subjected to *DpnI* (New England Biolabs Inc.) digestion followed by transformation. Positive clones were selected on a suitable antibiotic (kanamycin and ampicillin, respectively) containing plates and checked by the consequent release of correct-sized fragments following double digestion with *NdeI* and *BamHI* (New England Biolabs Inc.). All the clones were confirmed by sequencing.

The list of constructs and the primers used in this study are tabulated in Supplementary Table S3.

### Protein Expression

The constructs were transformed in BL21AI strains of *E. coli*. The cultures were grown in respective antibiotic-containing media (1X LB with 0.1 mg/ml ampicillin or 0.05 mg/ml kanamycin) and were kept in shaking conditions at 37°C. The cultures were induced with 0.2% L-Arabinose once it reached the exponential phase of growth (between 0.6-0.8 OD). The cultures were incubated overnight at 18°C. The samples were subjected to 15% SDS-PAGE to observe the overexpressed protein band of interest. Further, the selenomethionine-labeled protein was expressed in minimal media supplemented with selenomethionine using feedback inhibition method and was consequently purified.

### Protein purification

Purifications of MglA and MglB were performed as described in reference ^21^

#### His-tagged affinity purification

As a first step, the harvested cells were resuspended in the lysis buffer (50 mM Tris pH 8.0, 200 mM NaCl, and 10% glycerol) at 4°C. Consequently, the samples were centrifuged at 39,000 g for 45 minutes. The lysate was loaded on a 5 ml HisTrap HP (GE Healthcare) column. The loading buffer was 200 mM NaCl, 50 mM Tris pH 8.0, and the elution buffer was 200 mM NaCl, 50 mM Tris pH 8.0, and 500 mM Imidazole. Protein was eluted using a stepwise gradient of 2%, 5%, 10%, 20%, 50%, and 100% elution buffer.

#### Size Exclusion chromatography

For the RomX constructs, this step was performed after Ni-NTA affinity chromatography. The column was equilibrated with 50 mM NaCl, 50 mM Tris pH 8.0 (A50) buffer. For analytical runs, approximately 2-3 mg/ml, 200 µL of protein was injected into the Superdex 75 or Superdex200 size exclusion column (GE Healthcare) (volume less than 900 µL for preparative runs). UV absorbance at 280 nm was observed to monitor the elution of the protein. The respective fractions were pooled, concentrated, and stored.

#### Anion Exchange (MonoQ)

For RomR constructs, to improve the purity, this step was performed after Ni-NTA affinity chromatography. Hence, it was injected in MonoQ 4.6/100 PE (GE Healthcare). Buffers used for binding and elution were Buffer A (50 mM Tris [pH 8.0], 50 mM NaCl) and Buffer B (50 mM Tris [pH 8.0], 1 M NaCl), respectively. A linear gradient of Buffer A ranging from 0% to 50% Buffer B over 20 column volumes was injected, and the fractions containing the protein were pooled and concentrated.

### Crystallization and structure solution

To determine the experimental structure of RomX, constructs with both C-terminal and N-terminal hexahistidine tags, were subjected to crystallization screens. Nearly 500 conditions of commercially available screens (Molecular dimensions, Hampton Research, Jena Bioscience, Rigaku) were screened using Mosquito crystallization robotic system, with drop sizes of 100 nl of protein at a concentration of 15 mg/ml and 100 nl of crystallization condition, in 96-well sitting drop plates (MRC plate, SWISS-SCI) There were multiple hits obtained for C-His-RomX but none for N-His-RomX. This shows that the position of the hexahistidine tag might affect crystal packing and hence, consequently, affect the possibility of better crystallization of the protein. After multiple rounds of optimization, the condition that gave the best reproducible crystals was with 25% PEG MME 550, 5 mM MgCl_2_, and 50 mM HEPES Na, pH 7 using the hanging drop method. The crystals were long, needle-shaped. Since RomX was found to be a novel protein with no known structural homologs, we had to employ anomalous diffraction techniques using selenomethionine-labeled protein to determine the phase of the diffraction data to solve the crystal structure (Supplementary Fig. 1E). The protein was crystallized in the same aforementioned condition. The anomalous scattering data was obtained and an overall resolution of 1.9 Å was obtained for the best crystal.

Diffraction data was collected from Diamond Light Source, UK. RomX anomalous data were collected at a wavelength of 0.9793 with *f’*=8.03 and *f’’*=3.85. Data was reduced and scaled using iMOSFLM and AIMLESS, respectively, using the CCP4i2 package. Autosol and Autobuild wizards of PHENIX were used to build the model which was finally refined to satisfactory *R_work_* and *R_free_*values using iterative refinement cycles using Coot. The refinement statistics for each structure were tabulated to validate the quality of the refined data. We also obtained electron density corresponding to ethylene glycol which was used as a cryoprotectant for the diffracted crystal.

### SEC-MALS

Size exclusion chromatography coupled with Multi-angle light scattering (SEC-MALS) experiments enabled accurate molar mass estimation. The Superdex 75/200 Increase 10/300 GL column was used for SEC, which was connected to an Agilent HPLC unit with an 18-angle light scattering detector (Wyatt Dawn HELIOS II) and a refractive index detector (Wyatt Optilab T-rEX). The experiments were performed at room temperature. The column was equilibrated with A50 (50 mM NaCl, 50 mM Tris pH 8.0) buffer at 0.4 ml/min. BSA at 2 mg/ml was used to calibrate the system. The purified protein/complexes (approximately 5 mg/ml, 110 µL) were loaded to estimate the molecular weight of the eluted peaks. The Zimm model implemented in ASTRA software was used for the curve fitting and estimation of molecular weights. GraphPad Prism was used to average molar mass from fitted plots.

### Isothermal Titration calorimetry

The proteins (MglA, RomR-REC and RomX) used for interaction studies were dialyzed into 20 mM phosphate buffer (K_2_HPO_4_+KH_2_PO_4_), pH 8, overnight and the exact concentrations were estimated using Bradford assay. For MglA and RomX runs, 2 mM MgCl_2_ and 5 mM of nucleotide (GDP/GTP) were additionally supplemented in the reaction. ITC measurements were carried out by titrating 300 μM of RomX (N-terminal hexahistidine tagged construct) into 20 μM of RomR-REC in the sample cell using the MicroCal PEAQ-ITC, using a stirring speed of 750 rpm. 19 injections, 2 μl each were made. Similarly for MglA-RomX runs, measurements were carried out by titrating 1 mM of RomX (N-terminal hexahistidine tagged construct) into 50 μM of MglA in the sample cell using a stirring speed of 750 rpm. 19 injections, 1 μl each were made. Data for each experiment was modelled using a one-site binding model provided in the MicroCal PEAQ-ITC analysis software with fitted offset control. Error in K_d_, N, ΔG, ΔH and -TΔS was calculated as the standard error of the mean using multiple replicates.

### Tryptophan Fluorescence Assay

The experiment was performed in 20 mM phosphate buffer (K_2_HPO_4_+KH_2_PO_4_), pH=8. The sample volume was 200 μl in a quartz cuvette (10 × 2 mm path length) and excitation and emission slit widths of 2 nm. The excitation spectra were recorded between 200-300 nm. The emission spectra were recorded at the excitation wavelength of 287 nm and the emission spectra were recorded between 300-450 nm. 10 μM RomX was added to the buffer and the spectra were recorded. Further, it was titrated with 1-20 μM RomR in steps of 1 μM each. The emission maxima (□max) and the corresponding fluorescence intensity (in CPS, counts per second) were recorded and plotted against the concentration of RomR to estimate binding.

### CD Spectroscopy

Far-UV CD spectra were collected using a Jasco J-815 spectropolarimeter with a protein concentration of 10 μM in 20 mM phosphate buffer (K_2_HPO_4_+KH_2_PO_4_), pH=8, in a 1 mm cuvette, using a scan speed of 50 nm/min, a digital integration time of 2 s, and a bandwidth of 1 nm. Spectra was collected between 200 to 250 nm under permissive high-tension voltage. Each spectrum was averaged for over 30 scans. Spectra were deconvoluted and analyzed using BeStSel^60^.

### NADH-coupled GTP hydrolysis assay

NADH-coupled enzymatic assay ^61^, similar to the protocol used in reference ^21^was used to measure GTP hydrolysis activity. A master mix was prepared in A50 buffer (50 mM NaCl, Tris pH 8.0) containing 600 µM NADH, 1 mM phosphoenol pyruvate, 5 mM MgCl_2_, 1 mM GTP and pyruvate kinase and lactate dehydrogenase enzyme mix (∼25 U/ml). All components were mixed to a 200 µl reaction volume and added to a Corning 96 well flat bottom plate. The reactions were initiated by adding purified protein complexes in respective ratios to a final concentration of 10 μM MglA. A decrease in NADH absorbance was measured at 340 nm using a Varioskan Flash (4.00.53) multimode plate reader. The absorbance was measured every 20 s for 7200 s. The initial time point and absorbance of buffer components were subtracted from all readings. NADH absorbance was converted to GDP produced using a slope obtained from a standard curve containing known concentrations of NADH. GraphPad Prism was used for data analysis and plotting the *k_obs_*values.

### Nucleotide exchange assay

The kinetic measurements were performed on Fluoromax-4 (Horiba), where the intensity of fluorescence emission by *mant-*labeled GDP/GMPPNP (Jena Bioscience, Germany) at 440 nm was monitored after the excitation at 360 nm (protocol similar to the one used in^21^). The sample volume was 200 μl in a quartz cuvette (10 × 2 mm path length) and excitation and emission slit widths of 2 nm. *mant*-GDP/GMPPNP (final concentration 800 nM) was present in buffer A50 (50 mM Tris pH 8.0, 50 mM NaCl, 5 mM MgCl_2_). The protein mix, i.e., a final concentration of 3 μM of the respective protein complexes, was added in the cuvette at 400 seconds after stabilization of the signal from only *mant*-nucleotide. Consequently, the fluorescence was recorded for 1,400 seconds, and the increase in intensity reflected the nucleotide-binding kinetics. At 1,800 seconds, the *mant-*labeled nucleotide was competed out with an excess of unlabeled GDP/GTP (final concentration 500 μM), resulting in the release of *mant*-labeled nucleotide from the protein which manifested as a decrease of fluorescence intensity. For plotting the relative intensities from the measurements, each value was divided by the average of the first 200 readings (400 seconds). These accumulations and decay reactions were fitted to exponential binding equations as given below in GraphPad Prism to estimate the *k_on_*and *k_off_* values.

P + N_1_ → PN_1_ + N_2_ → PN_2_ + N_1_, where P represents protein, N_1_ is the labelled nucleotide, N_2_ is the unlabelled nucleotide, and PN denotes the protein-nucleotide complex.

For estimation of *k_on_*, PN_t_ = PN_max_ (1 − e^−kt^).

For estimation of *k_off_*, PN_t_ = PN_max_ − PN_min_ (e^−kt^) + PN_min_

Here, PN_t_ represents the amount of the complex at time t, PN_max_ is the maximum amount of the complex upon association, and PN_min_ is the minimum amount of the complex after dissociation.

### AlphaFold model prediction

Protein structure predictions for the MglA/RomX, RomR/RomX and MglA/RomR/X complexes were performed using AlphaFold v2.0^49^. The amino acid sequences of MglA, RomX and RomR were obtained from the UniProt database and merged into a single input file for complex structure prediction. AlphaFold was run with the multimer model setting, which is optimized for predicting protein-protein interactions. Five structural models were generated for each of the complexes. The predicted models were ranked based on the Predicted Local Distance Difference Test (pLDDT) score, which measures the confidence in the predicted position of each residue. Additionally, the Predicted Aligned Error (PAE) score was used to check the accuracy of the relative position of the two proteins in the complex. The best model was selected based on a combination of high pLDDT scores (>90) and PAE scores indicating stable interactions.

### Bioinformatics

Preliminarily, **t**he homologs of RomX were searched using the BLAST tool^62^. The secondary structure predictions of RomX and RomR were done using the PSIPRED web-server^63^.

The structure of RomX was given as input to the DALI server to perform a structure-based search against the Protein Data Bank database^40^. It returned several hits for proteins with one or more chains in the same fold as RomX (coiled-coil helices) from within the PDB. The top 50 structures in the DALI output were analyzed and based on the RMSD values obtained, a total of 19 structures were shortlisted for further analysis (Supplementary Table S1).

For the Foldseek search, the structure of RomX was given as input to search against AFDB Proteome, AFDB SwissProt, AFDB50, CATH50 MGNIFY_ESM30, and PDB100 databases^44,64^. Most of the hits were obtained from the AFDB50 database. Out of 471 hits, 169 proteins were characterized and 302 were of uncharacterized proteins. 52 hits contained response regulatory proteins from bacteria, 47 contained the Zinc finger domain, whereas the remaining folds had 10 or less number of hits. 26 hits were categorized as proteins of miscellaneous folds as they had a single unique unrelated fold for our study. The list of relevant domains and the corresponding AlphaFold ID is tabulated in Supplementary Table S2.

For sequence alignments of RomX and RomR-REC, NCBI BLASTp was used. Myxococcales (taxid:29) were excluded from the search. The hits were sorted according to the decreasing E-value. Alignments were generated using the top 500 hits and the conservation was represented using WebLogo^65^.

## Abbreviations

GEF: Gunanine nucleotide exchange factor
GAP: GTPase Activating Protein
*mant*: 2’/3’-O-(N-methyl anthraniloyl
GDP: guanosine diphosphate
GTP: guanosine 5’-tri-phosphate
PDB: Protein Data Bank
RomR: Regulator of motility response regulator
MglA: Mutual gliding protein A
MglB: Mutual gliding protein B
NADH: nicotinamide adenine dinucleotide
pLDDT: Predicted local distance difference test
ITC: Isothermal titration calorimetry
DALI: Distance Alignment matrix
REC: Receiver domain
SAD: Single wavelength anomalous diffraction

## Acknowledgements

The study was funded by the CEFIPRA Collaborative Research Grant (CSRP 5803-1) and supported by common instrumentation facilities at IISER Pune. SC acknowledges IISER Pune, Infosys Foundation, Biochemical Society, UK, DBT (DBT/CTEP/02/20220753076), and Indo-German Science & Technology Centre (IGSTC) for fellowships. The synchrotron facility at Diamond Light Source, UK (I03 beamline), and the macromolecular crystallography facility at the National Centre for Cell Science, Pune, are acknowledged for RomX data collection and crystal screening, respectively. Elettra Sincrotrone Trieste (XRD2; access facilitated by the Department of Science and Technology, Government of India) and the European Synchrotron Radiation Facility (ESRF; ID30A-3, ID23-1; access facilitated by Department of Biotechnology) are acknowledged for preliminary crystal screening for RomR-REC and RomR-REC/RomX complexes. Kushan Lahiri and Ranjana Nataraj are acknowledged for initiating the RomX fold search analyses and preliminary RomR-X docking studies. Dr Radha Chauhan and Jyotsna Singh from the National Centre for Cell Science, Pune, are acknowledged for their assistance in using the SEC-MALS facility. Suman Pal, Preeti Kumari and Prof. Jayant Udgaonkar are acknowledged for their assistance with CD spectroscopy experiments and access to the facility. Prof. Tam Mignot from CNRS, Marseille is acknowledged for providing the RomR plasmid from which the REC domain constructs were cloned.

## Author Contributions

SC: Designed and performed all the experiments and wrote the manuscript

PG: Conceptualized the study, supervised the experiments and analyses, and reviewed and edited the manuscript

## CrediT authorship contribution statement

SC: Methodology; Investigation; writing - original draft

PG: Conceptualization; Methodology; Funding acquisition; Project administration; Supervision; Writing - review and editing

## Declaration of competing interest

The authors declare no conflict of interest.

## Data Availability

The atomic coordinates and structure factors for RomX have been deposited in the Protein Data Bank (PDB) under the accession code 8ZNA.

## Notes

### Competing Interest Statement

The authors have declared no competing interest.

### Summary of Updates

A few updates on latest references and some new control experiments have been included.

