## Supplementary Figures and Tables for "RomX, a novel prokaryotic regulator, links the response receiver domain of RomR with GTP-bound MglA for establishing *Myxococcus xanthus* polarity"

### Supplementary information

**Table S1: List of structures containing a GTPase domain shortlisted from top 50 hits of DALI search using RomX structure as query**

| Chain | Name | Organism | Ligands bound | No. of residues | %id | RMSD with RomX |
|---|---|---|---|---|---|---|
| 6B9F-A | Atlastin-1 | *Homo sapiens* | GDP, ALF, GOL, MG | 411 | 13 | 0.816Å |
| 6B9D-A | Atlastin-1 | *Homo sapiens* | GDP, GOL, MG | 379 | 14 | 0.707Å |
| 4IDN-A | Atlastin-1 | *Homo sapiens* | GNP, MG | 417 | 14 | 0.755Å |
| 5CB2-A | Protein Sey1 | *Candida albicans* SC5314 | GNP. MG | 656 | 10 | 1.140Å |
| 5CA9-A | Protein Sey1 | *Candida albicans* SC5314 | GDP, ALF, MG | 656 | 14 | 0.778Å |
| 5CA8-A | Protein Sey1 | *Candida albicans* SC5314 | GDP, MG | 636 | 15 | 0.809Å |

**Table S2: List of hits from AFDB50 containing RomX fold with the respective domains and AlphaFold IDs**

| | |
|---|---|
| Response receiver domain protein | AF-A0A3D5Y9Q6,AF-A0A1F9CMP0,AF-A0A2V7W0H0,AF-A0A3B1DEH3,AF-A0A1F8ZLW2,AF-A0A2U3QHU4,AF-A0A1F5VDX1,AF-A0A1G1GZR7,AF-A0A7C2NLE3,AF-A0A6V8N7U0,AF-A0A1G1HLX1,AF-A0A1G1HZY9,AF-A0A2D6BR13,AF-A0A1F9PW42,AF-A0A2N6E6P0,AF-A0A2E9GUJ2,AF-A0A523JKP6,AF-A0A2V2RPA9,AF-A0A1E7IMD8,AF-A0A2E5YQE1,AF-A0A2E3NGZ3,AF-A0A2J6J6V7,AF-A0A2N2FU07,AF-A0A661PQ28,AF-A0A1X0YEX3,AF-A0A2D7HVL5,AF-A0A2H0YIM6,AF-A0A1Q7HYQ9,AF-A0A1I1T9F8,AF-A0A1Q7RSH3,AF-A0A7C4DI73,AF-A0A7C3L1W6,AF-A0A7C5YSD1,AF-A0A7X6ICB5,AF-A0A524GGA8,AF-A0A7C7RZZ4,AF-A0A523JXG4,AF-A0A533SEV2,AF-A0A533SD32,AF-A0A7C0WKH2,AF-A0A3M1Y533,AF-A0A7C8DJY7,AF-A0A7C7YCJ2,AF-A0A7C7YHP4,AF-A0A3M1KM66,AF-A0A7C7YGE4,AF-A0A3M1G3L1,AF-A0A523DUB7,AF-A0A3M1E8L1,AF-A0A0C2CZ80,AF-A0A2H5W6W0 |
| Zn finger/ thioredoxin put. domain protein | AF-A0A660S8H5,AF-A0A538T2R0,AF-A0A7V3ZSY5,AF-A0A538T8F1,AF-A0A496UVP3,AF-A0A2G6G5P2,AF-A0A523Z6Y8,AF-A0A2M7YVX7,AF-A0A1V4QEG8,AF-A0A538TMC5,AF-A0A124G0L7,AF-A0A7C6AFL9,AF-A0A7C4U9K4,AF-A0A660SHX7,AF-A0A7C4TDL3,AF-A0A3M2BRZ5,AF-A0A1Y1RII6,AF-A0A6C1NT35,AF-A0A7X5WCK7,AF-A0A2W4U9S6,AF-A0A523XHU7,AF-A0A2V7MQD7,AF-A0A2V7JF24,AF-A0A7X7SI64,AF-A0A660SI27,AF-A0A7V2AUQ1,AF-A0A651DKE9,AF-A0A2V7PVT9,AF-A0A6M4IQK4,AF-A0A2V7R5T7,AF-A0A2V7SVV4,AF-A0A1F5V4N6,AF-A0A2V7M8N4,AF-A0A2G6LXI9,AF-A0A2V7RTR3,AF-A0A7Y2EA27,AF-A0A3B8UYI4,AF-A0A6B1FLZ9,AF-A0A1V6K5Z6,AF-A0A2V8TER6,AF-A0A0Q1AE57,AF-A0A1F5RPX1,AF-A0A538U3Z4,AF-A0A538UB26,AF-A0A538SF55,AF-A0A538SKG2,AF-A0A841H474 |
| GAF domain protein | AF-A0A3M1PV69,AF-A0A2V8DT00,AF-A0A7V1UM56,AF-A0A7V9UV04,AF-A0A7T9FPN5,AF-A0A7W0UY03,AF-A0A2V8IZQ7,AF-A0A7Y5U5S2,AF-A0A2V8I2C3,AF-A0A522PYD8 |
| DUF domain protein | AF-A0A521UUF1,AF-A0A549TBY5,AF-G4QJS9,AF-A0A426T061,AF-A0A3B8Y644,AF-A0A5M4D8A0,AF-A0A4Y2F1X1,AF-A0A4Y2MRZ8,AF-A0A4Y2NIX5,AF-A0A4Y1ZK84 |
| Zinc-ribbon domain protein | AF-A0A7Y7NPC9,AF-A0A838W9R4,AF-A0A7V9B8N1,AF-A0A7W1BHI6,AF-A0A7V9TMC9,AF-A0A7W0W6S0,AF-A0A7V9U1W3 |
| GAT domain protein | AF-A0A1F5LPA7,AF-A0A5N5DIJ6,AF-A0A1S8B605,AF-C9SBW5, AF-A0A0B1P008 |
| Sigma70 factors | AF-A0A7V1EPE6,AF-A0A7Y3EF57,AF-A0A5S4VMB1 |
| Histidine kinase | AF-A0A3B9VE56,AF-A0A2N6G7Z6 |
| Chromosome segregation ATPases-like protein | AF-Q1IN57,AF-A0A136JSP0 |
| **Ribosome-associated GTPase EngA** | AF-A0A085FXZ6 |
| **eIF-2B GDP-GTP exchange factor subunit alpha** | AF-A0A420HVJ4 |
| AAA domain protein | AF-S0FI48 |
| Chemotaxis protein CheY | AF-A0A0D5N4I1 |

**Table S2: List of constructs and primers used in the study**

| Construct | Primers |
|---|---|
| *pHis17 romX H6*<br>$Amp^R$ | RomX-F<br>5'GTTTAACTTTAAGAAGGAGATATACATATGACGGACGAGGAAAAGGTC3'<br>RomX-H6-R<br>5'GCTTTTAATGATGATGATGATGATGGGATCCCCAGATCTTCGACTTCAC3' |
| *pHis17 H6 romX*<br>$Amp^R$ | RomX-H6-F<br>5'ATATACATATGCGTGGCCATCATCATCATCATCATGGAGGAACGGACGAGGAAAAGGTC3'<br>RomX-R<br>5'TTTAATGATGATGATGATGATGGGATCCTTACCAGATCTTCGACTTCAC3' |
| *pHis17 H6 romX L75A,R76A*<br>$Amp^R$ | RomX-F<br>5'GTTTAACTTTAAGAAGGAGATATACATATGACGGACGAGGAAAAGGTC3'<br>RomX-L75A R76A-H6-R<br>5'CGACTTCACATGCGCCTTCGACGCCGCGACGATGTCGTTGATCGC3' |
| *pHis17 H6 romX D19A*<br>$Amp^R$ | RomX-D19-F<br>CTCGCCCGTGCGATTGCCTCGGCTATCTCGCTCTACAACGAGCAGAAG<br>RomX-R<br>5'TTTAATGATGATGATGATGATGGGATCCTTACCAGATCTTCGACTTCAC3' |
| *pHis17 H6 romX R11,14A*<br>$Amp^R$ | RomX-RR-F<br>CTCGCCCGTGCGATTGCCTCGGCTATCTCGCTCTACAACGAGCAGAAG<br>RomX-R<br>5'TTTAATGATGATGATGATGATGGGATCCTTACCAGATCTTCGACTTCAC3' |
| *pHis17 romR REC116 H6*<br>$Kan^R$ | RomR-F<br>5'GTTTAACTTTAAGAAGGAGATATACATATGCCCAAGAATCTGCTG3'<br>RomR-116-H6-R<br>5'GCTTTTAATGATGATGATGATGATGGGATCCCACCTTGTCGAGCAGCACCTG3' |
| *pHis17 romR REC125 H6*<br>$Kan^R$ | RomR-F<br>5'GTTTAACTTTAAGAAGGAGATATACATATGCCCAAGAATCTGCTG3'<br>RomR-125-H6-R<br>5'GCTTTTAATGATGATGATGATGATGGGATCCGTTGGACTTCTGGCCGACCAGCGCCTT3' |
| *pHis17 romR REC141 H6*<br>$Kan^R$ | RomR-F<br>5'GTTTAACTTTAAGAAGGAGATATACATATGCCCAAGAATCTGCTG3'<br>RomR-125-H6-R<br>5'GCTTTTAATGATGATGATGATGATGGGATCCCTGAGGAGCCGCGTGGCGCACCTGCGT3' |
| *pHis17 romR 331Ct H6*<br>$Kan^R$ | RomR-331Ct-F<br>5'GTTTAACTTTAAGAAGGAGATATACATATGCCGTCCATCAGCATCGAGGACTCGCTG3'<br>RomR-H6-R<br>5'GCTTTTAATGATGATGATGATGATGGGATCCGTGCTGGGTCTCTC3' |
| *pHis17 romR REC E108A H6*<br>$Kan^R$ | RomR-F<br>5'GTTTAACTTTAAGAAGGAGATATACATATGCCCAAGAATCTGCTG3'<br>RomR-125-H6-R<br>5'CTTCACCTTGTCGAGCAGCACCTGGCTCGCGAAGGGCTTGGTGAC3' |

### Supplementary Figure S1

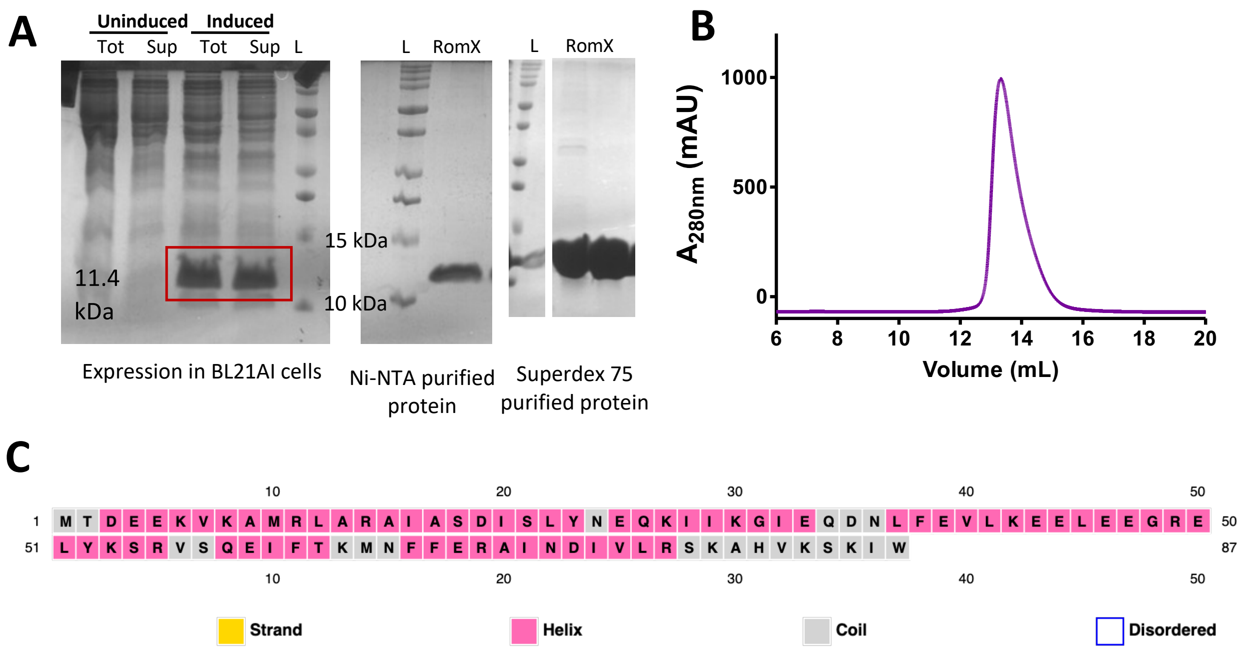


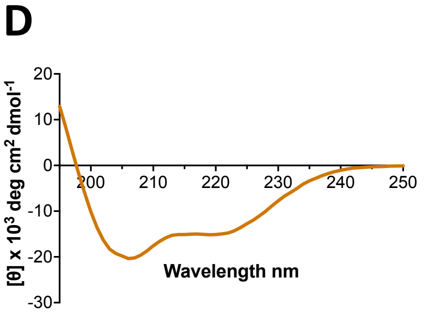


| Secondary Structure | Content (%) |
|---|---|
| α helix | **50.4** |
| Antiparallel β strands | 7.8 |
| Parallel β strands | 0.8 |
| Turn | 0.0 |
| Others | 40.9 |

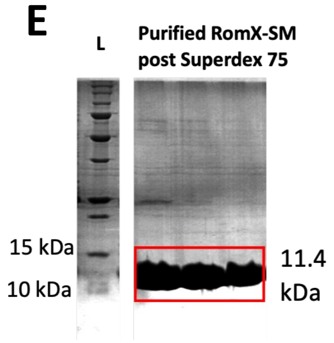


**Figure S1: Structural characterization of RomX**

A: SDS PAGE for expression of N-His RomX. The expressed protein band at 11.4 kDa in induced cultures is highlighted in red. The right panels are SDS PAGE for Ni-NTA purified and Superdex 75 purified protein.

B: Superdex 75 analytical profile for hexahistidine tagged RomX. The protein elutes at 13.8 ml.

C: Secondary structure prediction of RomX using PSIPRED shows it to be composed primarily of alpha helices.

D. The CD spectrum of the RomX protein predicts abundance of alpha helices in the protein structure as per the BeStSel deconvolution shown in right.

E. SDS PAGE for purification of C-His RomX grown in minimal media supplemented with Selenomethionine. The protein was purified using Ni-NTA followed by Superdex 75. The profiles were the same as earlier purifications.

### Supplementary Figure S2

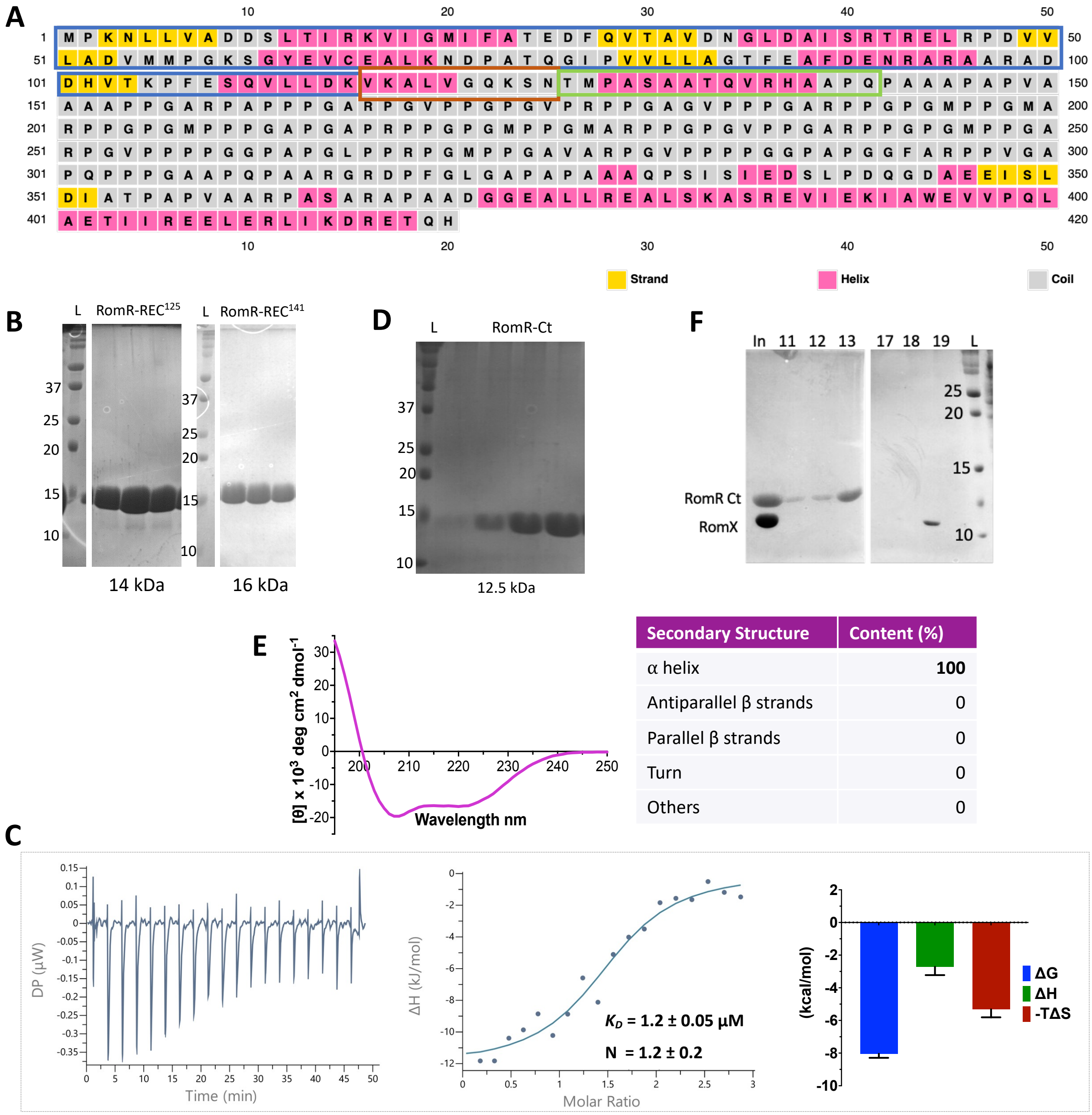


| Secondary Structure | Content (%) |
|---|---|
| α helix | 100 |
| Antiparallel β strands | 0 |
| Parallel β strands | 0 |
| Turn | 0 |
| Others | 0 |

**Figure S2: Interaction of RomR domains with of RomX**

A: Secondary structure prediction of RomR using PSIPRED. The three different boundaries of the REC domain are highlighted in blue (116), orange (125) and green (141), respectively. The C-terminal domain is highlighted in magenta.

B, D: SDS PAGE showing purified proteins of the three REC domain constructs and C-terminal domain construct of RomR, respectively, after anion exchange chromatography based purification.

C: ITC thermogram showing the titration of RomX with RomR-REC$^{141}$. The left panel shows the raw data of exothermic heat pulses with time, the middle panel shows the corresponding differential binding curve fitted to a single site binding model and the right panel shows the thermodynamic signature plot detailing the ΔG, ΔH and -TΔS values corresponding to the same reaction (mean and standard errors are shown).

E: CD spectrum of the RomR-331Ct protein was fed on to BeStSel server to obtain the percentage of estimated secondary structure content. The fitting shows a 100% alpha-helical content.

F: SDS-PAGE corresponding to analytical SEC run with RomR-Ct with RomX (Fig. 2E).

### Supplementary Figure S3

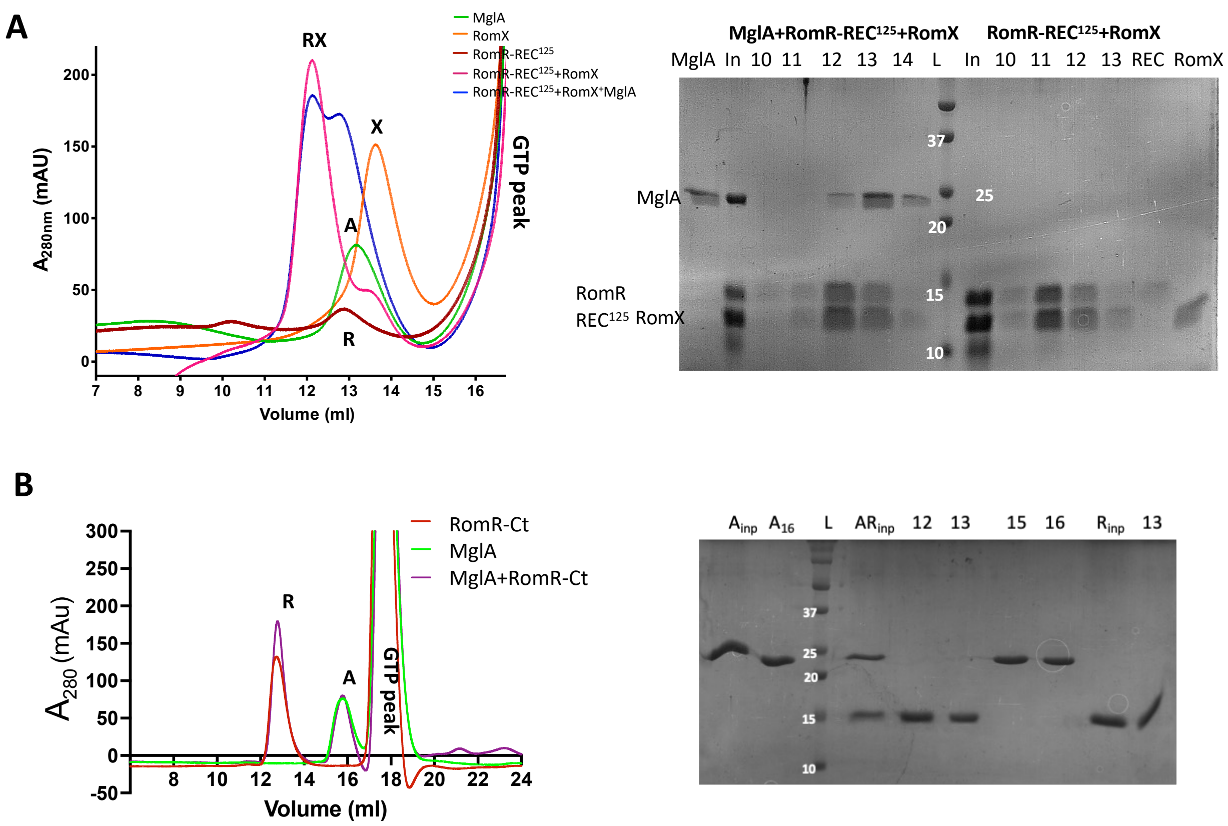


**Figure S3: Interaction of RomR-RomX complex with MglA**

A: Analytical Superdex 75 runs showing the interaction of RomR-REC$^{125}$ with RomX and MglA with GTP. The axis is cut to mask the high peaks of GTP elution at 18 ml, as shown in the figure. The right panel shows the corresponding SDS PAGE images of the respective runs. The lane numbers indicate the volumes of elution as per the chromatograms (In: Input, L: Ladder, molecular weights are labelled) and the proteins corresponding to the bands are labelled. The corresponding peaks in the chromatograms are also labelled.

MglA (green), RomX (orange), RomR-REC$^{125}$ (brick red), RomR-REC$^{125}$ + RomX (magenta) and MglA+ RomR-REC$^{125}$ + RomX (blue)

B: Analytical Superdex 200 runs showing the interaction of RomR-Ct with MglA with GTP. The axis is cut to mask the high peaks of GTP elution at 18 ml, as shown in the figure. The right panel show the corresponding SDS PAGE images of the respective runs. The lane numbers indicate the volumes of elution as per the chromatograms (In: Input, L: Ladder) and the proteins corresponding to the bands are labelled. The corresponding peaks in the chromatograms are also labelled.

MglA (green), RomR-Ct (brick red), and MglA+ RomR-Ct (purple)

### Supplementary Figure S4

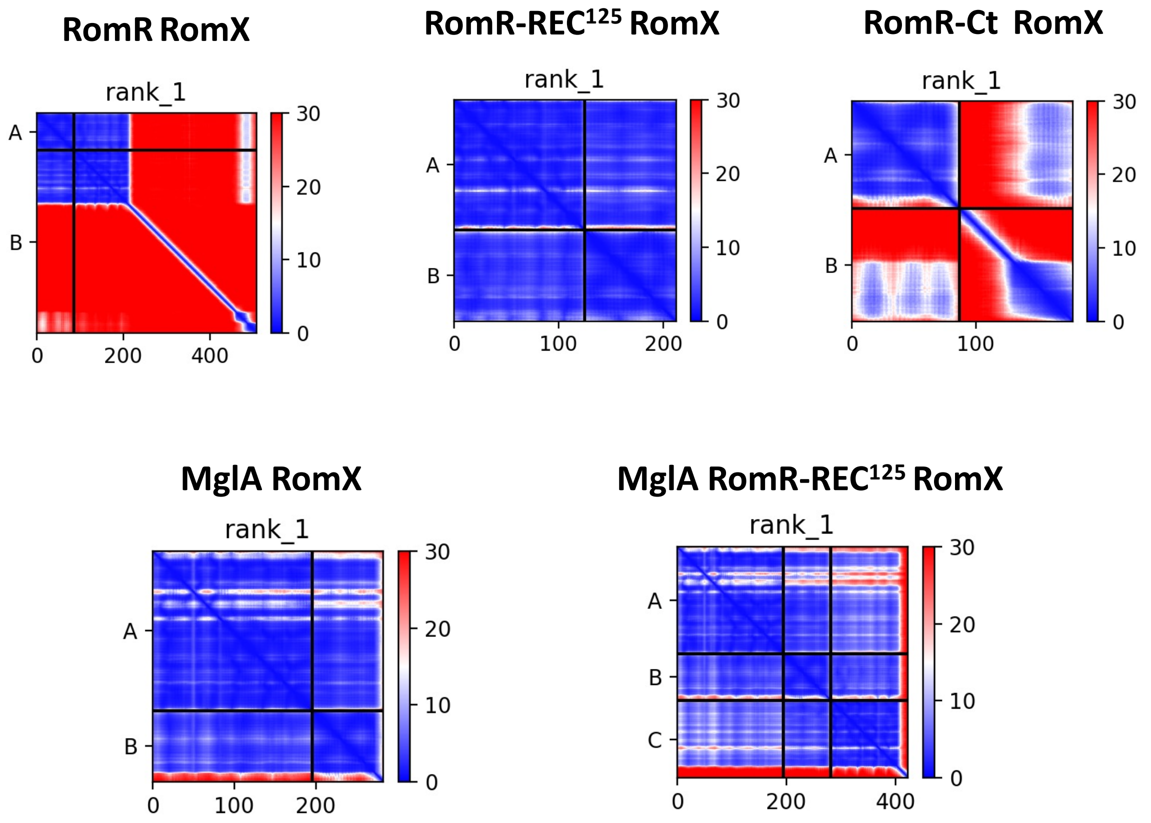


**Figure S4: PAE scores for the predicted AlphaFold models depicted in this study**

### Supplementary Figure S5

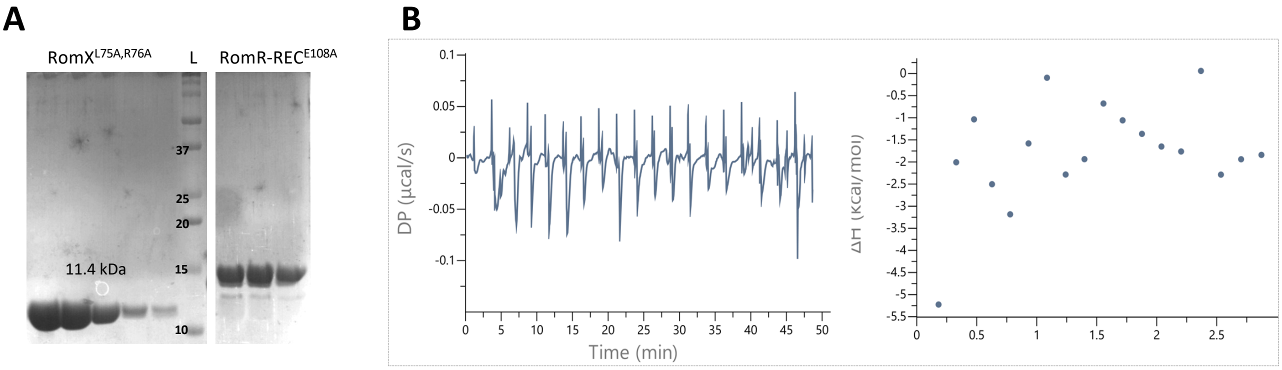


**Figure S5: Interaction interface between RomR-REC domain and RomX**

A: SDS-PAGE showing the purified protein for the RomX (L75A, R76A) and RomR-REC (E108A) interface mutants, post Superdex 75 and anion exchange, respectively.

B: ITC thermogram of a representative binding assay showing the titration of RomX$^{L75,R76A}$ with RomR-REC$^{E108A}$. The top panel shows the raw data of heat pulses with time. The lower panel shows the corresponding heat exchange, which could not be reliably fitted to a binding curve, indicating no binding.

### Supplementary Figure S6

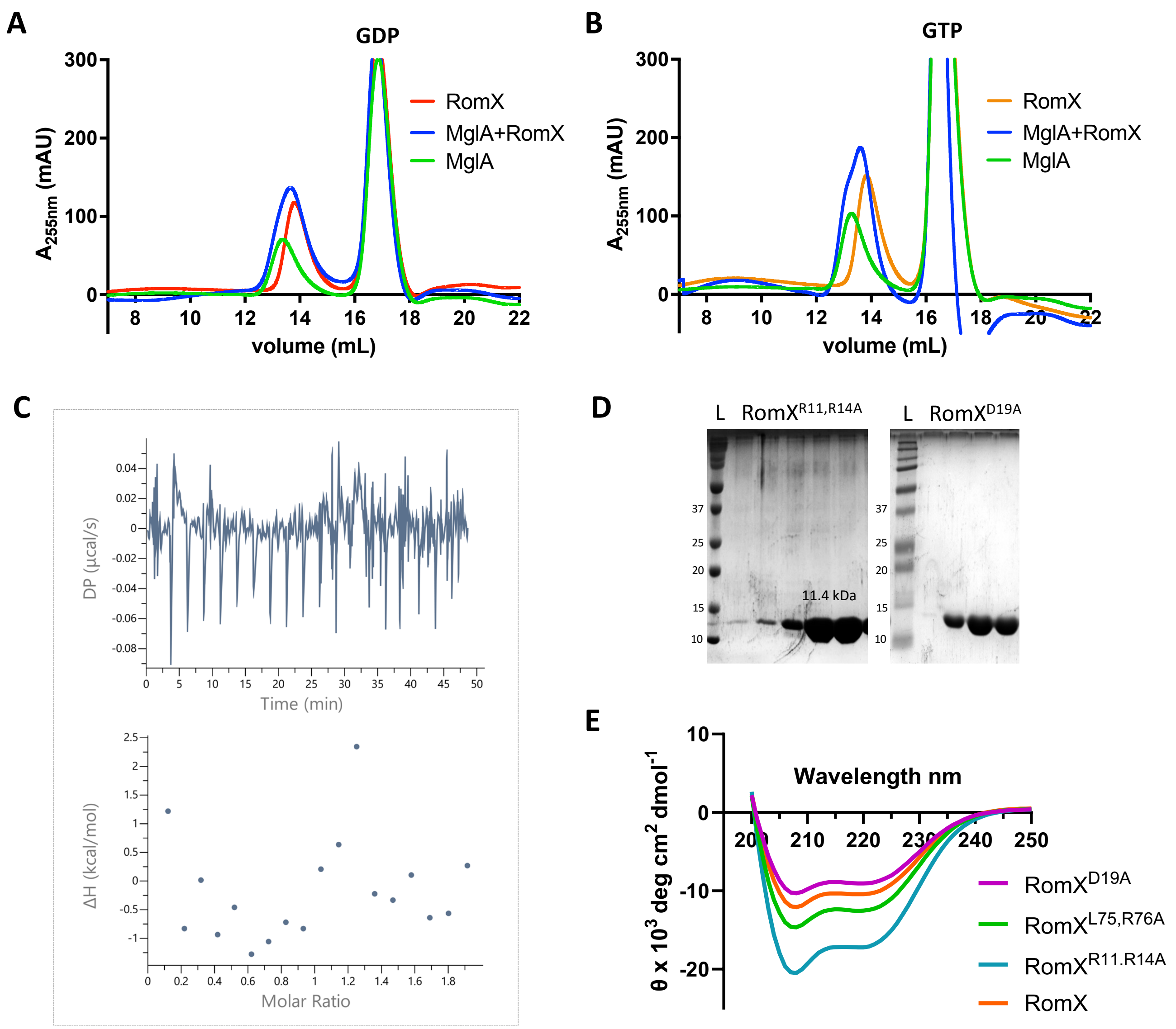


**Figure S6: Deciphering RomX-MglA interacting interface**

A, B: Analytical SEC using Superdex 75 column to detect binding of RomX to MglA in presence of GDP and GTP, respectively. No significant binding was observed as MglA (magenta) and RomX (blue) eluted at very close volumes between 13-14 ml and there was no obvious shift in the elution volume at the peak of the MglA+RomX (green).

C: ITC thermogram of a representative binding assay showing the titration of RomX with MglA in the presence of GDP and magnesium. The top panel show the raw data of heat pulses with time. The lower panel show the corresponding heat exchange, which could not be reliably fitted to a binding curve, indicating no binding.

D: SDS-PAGE showing the purified protein for the RomX mutants (R11,R14A and D19A, respectively) post Superdex 75.

E: The CD spectra of the RomX protein along with all the mutants of RomX used in this study ($RomX^{R11,R14A}$ in cyan, $RomX^{D19A}$ in magenta, $RomX^{L75,R76A}$ in green, and wild type RomX in orange.
